# Disease-specific eQTL screening reveals an anti-fibrotic effect of AGXT2 in nonalcoholic fatty liver disease

**DOI:** 10.1101/2021.03.22.436368

**Authors:** Taekyeong Yoo, Sae Kyung Joo, Hyo Jung Kim, Hyun Young Kim, Hyungtai Sim, Jieun Lee, Hee-Hoon Kim, Sunhee Jung, Youngha Lee, Oveis Jamialahmadi, Stefano Romeo, Won-Il Jeong, Geum-Sook Hwang, Keon Wook Kang, Jae Woo Kim, Won Kim, Murim Choi

## Abstract

**Background & Aims:** Nonalcoholic fatty liver disease (NAFLD) poses an impending clinical burden. Genome-wide association studies have revealed a limited contribution of genomic variants to the disease, requiring alternative but robust approaches to identify disease-associated variants and genes. We carried out a disease-specific expression quantitative trait loci (eQTL) screen to identify novel genetic factors that specifically act on NAFLD progression on the basis of genotype.

**Methods:** We recruited 125 Korean biopsy-proven NAFLD patients and healthy individuals and performed eQTL analyses using 21,272 transcripts and 3,234,941 genotyped and imputed SNPs. We then selected eQTLs that were detected only in the NAFLD group, but not in the control group (*i.e*., NAFLD-eQTLs). An additional cohort of 162 Korean NAFLD individuals was used for replication. The function of the selected eQTL toward NAFLD development was validated using HepG2, primary hepatocytes and NAFLD mouse models.

**Results:** The NAFLD-specific eQTL screening yielded 242 loci. Among them, *AGXT2*, encoding alanine-glyoxylate aminotransferase 2, displayed decreased expression in NAFLD patients homozygous for the non-reference allele of rs2291702, compared to no-NAFLD subjects with the same genotype (*P* = 4.79 × 10^−6^). This change was replicated in an additional 162 individuals, yielding a combined *P*-value of 8.05 × 10^−8^ from a total of 245 NAFLD patients and 48 controls.

Knockdown of *AGXT2* induced palmitate-overloaded hepatocyte death by increasing ER stress, and exacerbated NAFLD diet-induced liver fibrosis in mice. However, overexpression of AGXT2 reversely attenuated liver fibrosis and steatosis as well.

**Conclusions:** We implicate a new molecular role of AGXT2 in NAFLD. Our overall approach will serve as an efficient tool for uncovering novel genetic factors that contribute to liver steatosis and fibrosis in patients with NAFLD.

**Lay summary:** Elucidating causal genes for NAFLD has been challenging due to limited tissue availability and the polygenic nature of the disease. Using liver and blood samples from 125 biopsy-proven NAFLD and no-NAFLD Korean individuals and an additional 162 individuals for replication, we devised a new analytic method to identify causal genes. Among the candidates, we found that AGXT2-rs2291702 protects against liver fibrosis in a genotype-dependent manner with the potential for therapeutic interventions. Our approach enables the discovery of NAFLD causal genes that act on the basis of genotype.

## Introduction

Nonalcoholic fatty liver disease (NAFLD) is a growing burden that affects approximately a quarter of the world’s population and contributes to liver-related morbidity.[1,2] Defined as a condition in which excess liver fat exists in the absence of secondary causes of lipid accumulation or clinically significant alcohol intake, NAFLD includes a spectrum of liver diseases ranging from nonalcoholic fatty liver (NAFL) to nonalcoholic steatohepatitis (NASH).[1,3,4] Due to unmet needs in the prediction and early detection of NAFLD, there have been continuous efforts at clarifying its pathomechanism to facilitate the identification of novel therapeutic targets and biomarkers. However, no pharmacotherapy has yet been approved for NAFLD.[5,6]

Genome-wide association studies (GWAS) have revealed loci that confer risk for NAFLD.[1,4,7,8] However, these signals demonstrate modest effect sizes and account for only a minor fraction of the overall heritability of NAFLD, which is estimated to range at between 22–50%.[4,9] For complex traits, the development of a polygenic risk score (PRS) has led to promising results in risk prediction.[7] However, PRS evaluation of NAFLD has not yet generated robust results.[10,11] Mapping of expression quantitative trait loci (eQTLs) enables the identification of genetic variants that are associated with gene expression changes.[12,13] eQTL analysis has the advantage of providing interpretable molecular links between genetic variants and traits of interest.[14,15] These links also enable a substantial increase in statistical power, such that thousands of eQTLs can be detected even with just a sample of ~100 individuals.[14,16]

Alanine-glyoxylate aminotransferase 2 (AGXT2) is a mitochondrial aminotransferase, possessing multiple enzymatic activities on a wide array of substrates, including asymmetric dimethylarginine (ADMA) and 3-amino-isobutyrate (BAIB), and produces diverse metabolites including dimethylguanidino valeric acid (DMGV).[17] Knockout mouse and human-based studies have implicated *AGXT2* in endothelial dysfunction, hypertension, and chronic heart failure, as ADMA is a potent inhibitor of nitric oxide synthase (NOS).[18] GWAS of BAIB and DMGV levels in urine and serum indicated a strong association with *AGXT2* variants, highlighting its central role in regulating the levels of these molecules.[19–21] The gene is specifically expressed in the kidney and liver, and associated eQTLs have been identified in the liver tissues.[22] However, the molecular basis of its function in the liver remains elusive. Moreover, its pathophysiological role beyond the known enzymatic activity has not yet been clarified.

To circumvent existing technical limitations in understanding NAFLD, we performed eQTL mapping to identify genetic variants and their associated genes that confer susceptibility to NAFLD by collecting histologically confirmed liver tissue transcriptome and genotype data from 125 Korean individuals. We then developed a pipeline to select gene-eQTL pairs that are specifically active under the diseased state (hereafter referred to as “NAFLD-eQTLs”) and pinpointed a SNP in the *AGXT2* locus. With these efforts, we demonstrated that altered *AGXT2* expression might modulate the progression of liver fibrosis via ER stress-mediated hepatocellular death. Overall, our results highlight a new approach for evaluating and selecting NAFLD-eQTLs, which can lead to the identification of novel therapeutic targets for NAFLD in an individual-specific manner.

## Materials and Methods

### Subjects

This study was approved by the institutional review board of Seoul Metropolitan Government Boramae Medical Center. We constructed a prospective cohort from the ongoing Boramae nonalcoholic fatty liver disease (NAFLD) registry (NCT 02206841) as previously described.[23] See the Supplementary Methods for eligibility and diagnostic criteria used in the study. Participants of both discovery (*n* = 125) and replication (*n* = 162) cohorts consisted of Korean individuals (Fig. S1), aged 19–80, who visited Seoul Metropolitan Government Boramae Medical Center. All participants were informed of the study protocol and provided written and signed consent. NAFLD activity scoring and fibrosis staging were performed following the Kleiner classification, and categorized into no-NAFLD, NAFL, and NASH (Table 1, Table S1 and S2).[24] In the subsequent analyses, we considered no-NAFLD as control and NAFL and NASH as the NAFLD group.

**Table 1.** Summary statistics of the participants (*n* = 125).

|  | No-NAFLD<br>(*n* = 42) | NAFLD<br>(*n* = 83) | *P*-value |
| --- | --- | --- | --- |
| Age, years | 56.7 (12.3) | 53.3 (14.1) | 0.19 |
| Male, N (%) | 25 (59.5) | 38 (45.8) | 0.15 |
| BMI, kg/m<sup>2</sup> | 24.0 (3.4) | 28.1 (4.0) | 1.21 × 10<sup>-7</sup> |
| Fibrosis stage (0-4) | 0.26 (0.5) | 1.66 (1.0) | 4.45 × 10<sup>-19</sup> |
| Significant fibrosis (≥2) (%) | 1 (2.4) | 40 (48.2) | 2.56 × 10<sup>-7</sup> |
| NAFLD activity score | 0.43 (0.6) | 4.65 (1.3) | 3.60 × 10<sup>-49</sup> |
| SBP, mmHg | 131.0 (16.5) | 130.9 (18.5) | 0.98 |
| DBP, mmHg | 79.4 (11.2) | 79.9 (12.7) | 0.84 |
| HDL-cholesterol | 46.4 (10.4) | 44.7 (13.0) | 0.45 |
| Triglycerides, mg/dL | 123.7 (54.8) | 171.0 (88.0) | 3.72 × 10<sup>-4</sup> |
| AST, IU/L | 29.9 (18.7) | 67.5 (72.1) | 1.91 × 10<sup>-5</sup> |
| ALT, IU/L | 30.3 (24.8) | 80.5 (76.0) | 2.51 × 10<sup>-7</sup> |
| GGT, IU/L | 49.4 (53.8) | 66.7 (49.0) | 8.72 × 10<sup>-2</sup> |
| Albumin, g/dL | 4.1 (0.3) | 4.1 (0.3) | 0.44 |
| Platelet, ×10<sup>9</sup>/L | 228.7 (50.7) | 224.4 (64.0) | 0.68 |
| HOMA-IR | 2.7 (1.1) | 6.2 (4.8) | 9.11 × 10<sup>-9</sup> |
| Adipo-IR | 5.8 (4.5) | 13.8 (11.4) | 1.77 × 10<sup>-7</sup> |
| Diabetes, N (%) | 9 (21.4) | 41 (49.4) | 2.57 × 10<sup>-3</sup> |
| Hypertension, N (%) | 16 (38.1) | 31 (37.3) | 0.94 |

Values are given as mean (SD). *P*-values are from independent *t*-tests and $\chi^2$ tests
comparing between no-NAFLD and NAFLD groups.

### Transcriptome and genome data processing

Total RNA isolated from liver was used for RNA sequencing on a HiSeq2500 platform. Reads were mapped and quantified with the human genome (hg19/GRCh37) based on GENCODE v19. Differentially expressed genes (DEGs) were called using the DESeq2 packages[25] with correction for sample batches. Genes within certain criteria were used in eQTL analysis (see Supplementary Methods).

DNA was acquired from blood and genotyped using an Illumina Infinium OmniExpress-24 kit or Omni2.5-8 kit. Genotype data were processed with exclusion criteria, matched with RNA-seq data and imputed using 1000 Genome Project Phase 3 haplotypes. After a further filtering process, genotyped and imputed calls were used in eQTL analysis (see Supplementary Method).

### Cis-eQTL analysis

Genotype and gene expression data (21,272 genes and 3,234,941 genotyped and imputed SNPs) available from the 125 individuals were integrated for eQTL mapping using MatrixEQTL (version 2.2),[26] accounting for age and sex (or with body mass index (BMI) and homeostatic model assessment of insulin resistance (HOMA-IR)). MatrixEQTL performed a linear regression on the transformed residuals with the corresponding imputed genotypes under the additive model. An eQTL within 1 Mb of a gene transcription start site (TSS) was considered a *cis*-eQTL. False discovery rate (FDR) was used to adjust for multiple testing.

### Selecting NAFLD-specific eQTLs (NAFLD-eQTLs)

To choose NAFLD-eQTLs, we divided samples into no-NAFLD (*n* = 42) and NAFLD (*n* = 83) groups as described above and performed *cis*-eQTL calling separately. Among the eQTLs from the NAFLD group, calls that were also found in the no-NAFLD group with FDR-adjusted *P* < 0.05 were excluded. To define NAFLD-specific loci, we selected the most significant eSNP per eGene, then keeping only eQTLs with absolute β coefficient > 0.5 in NAFLD and absolute β fold change (FC) between NAFLD and no-NAFLD > 5. Finally, NAFLD-specific eGenes that overlapped with GTEx liver eGenes were excluded. GTEx participants free of NAFLD signatures (*n* = 79) were used. RNA-seq and genotyping data from GTEx release v7 (https://gtexportal.org) were downloaded from the Database of Genotypes and Phenotypes (dbGaP) under accession phs000424.v7.p2.[27]

### Agxt2 overexpression and knockdown in mouse model

Male C57BL/J mice were purchased from Japan SLC (Shizuoka, Japan). Six-week-old mice were fed for three weeks, either a normal chow-diet for a normal control model or a choline-deficient, L-amino acid-defined, high-fat diet (CDAHFD, Research Diets, New Brunswick, NJ) for a NAFLD model. Then, mice were injected with adenoviruses containing short hairpin RNA (shRNA) against *Agxt2* or mock, or adenoviruses expressing *Agxt2*-FLAG or GFP for knockdown and overexpression experiments, respectively. All procedures were performed under the standard protocols approved by the Committee on Animal Investigations of Yonsei University.

## Results

### Analysis of eQTLs

For transcriptome and genome-wide array analyses, liver biopsy and blood samples were acquired from 125 histologically-confirmed Korean individuals with varying metabolic and histological status. Using standardized pathological scores, we divided all samples into no-NAFLD (*n* = 42) and NAFLD (*n* = 83, including NAFL and NASH individuals) groups for eQTL analysis (Fig. 1A and Table 1 and S1). After quality assessment of the genotype and transcriptome data, 21,272 transcripts and 3,234,941 SNPs were subject to *cis*-eQTL (*i.e*., SNP-gene pair within 1 Mb of gene transcription start site (TSS)) mapping using an additive linear model. We identified 3,882 genes with *cis*-eQTLs at an FDR ≤ 5% (eGenes, Table S3), and 242,691 significant SNP-gene pairs from the set of 125 samples (“Liver-eQTLs”). We compared our liver eGenes to those from the GTEx database and found 40.1% of our eGenes (1,558/3,882) overlapped with GTEx liver eGenes, which was the largest proportion of overlap across all 48 GTEx tissues tested (Fig. S2A). Among the eQTL SNPs (eSNPs), 23.0% acted on multiple genes. As sample size may serve as a confounding factor for eQTL detection sensitivity, we compared our result to GTEx sets in terms of the correlation between sample numbers and eGene numbers. Our dataset fell within the range of GTEx correlations, suggesting that our data processing and eQTL calling were well-performed (Pearson’s correlation *R* = 0.94; *P* = 1.55 × 10^-24^; Fig. S2B). In addition, we detected 2,577 significant *trans*-eQTLs at FDR ≤ 5%, of which 646 (25.1%) were also eQTLs for nearby genes. Among the *trans*-eQTLs, 30.0% predicted the expression of multiple non-local genes. We limited our subsequent analyses on *cis*-eQTLs as they harbor stronger and more direct implications on target gene expression regulation.

**Figure 1.**
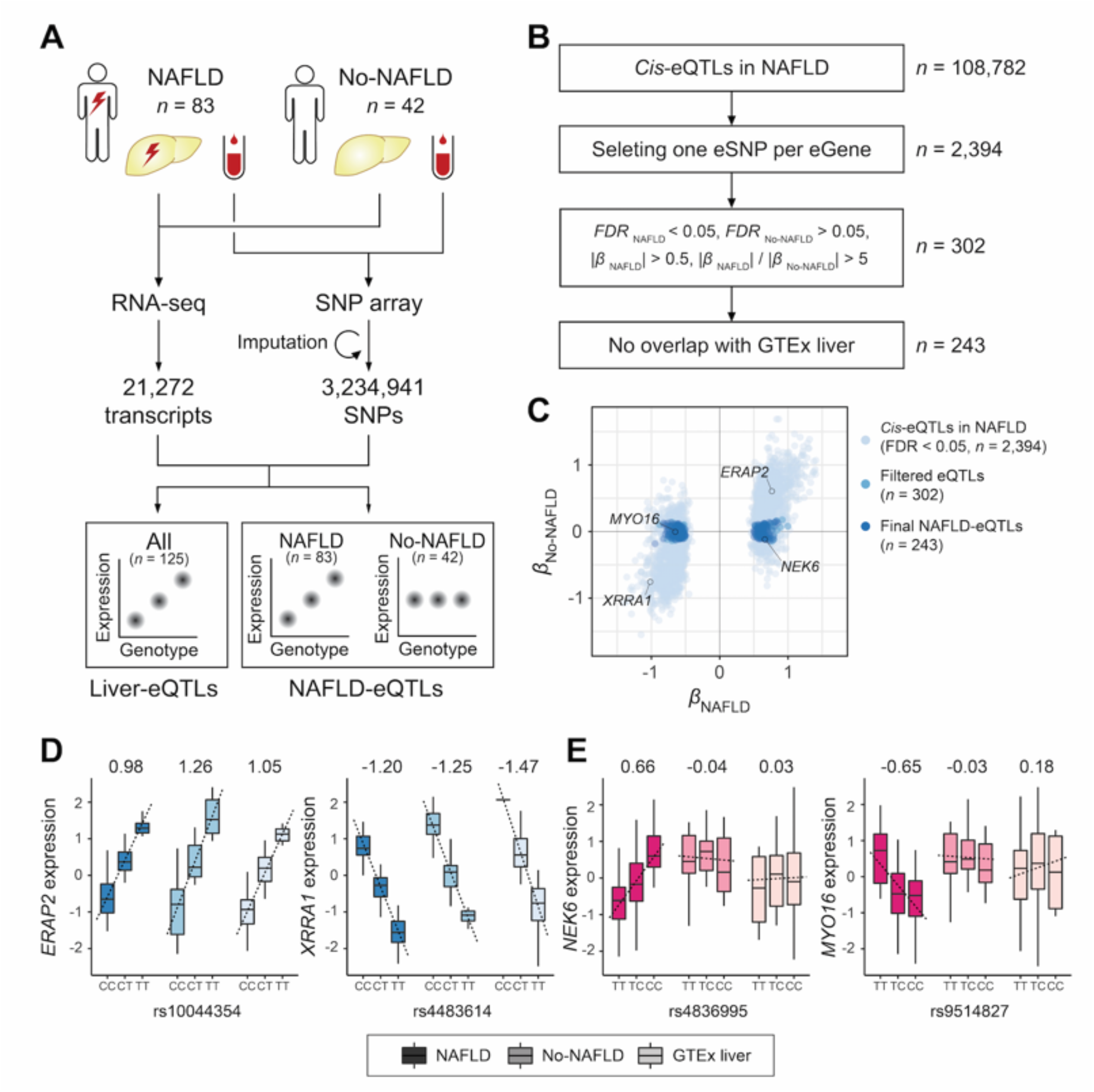
Screening NAFLD-eQTLs. (A) Study scheme. (B) NAFLD-eQTL calling process. (C) Scatterplot of β values depicting the eQTL calls during the filtering process described in (B). Genes shown in D-E are indicated. (D-E) Examples of liver-QTLs that show significant associations in all physiological conditions (D), and of NAFLD-eQTLs that are significant only in the NAFLD group (E). The numbers above each genotype class denote β values. Darkness of bars depicts different groups (NAFLD, no-NAFLD, and GTEx liver).

### Calling NAFLD-eQTLs

Under the hypothesis that *cis*-eQTLs may possess different activities under altered physiological status, such as NAFLD, we called eQTLs that are significantly associated in NAFLD patients but not in the no-NAFLD group (*i.e*., NAFLD-eQTLs). Using the same sets of SNPs and genes as for the liver-eQTL calling as described above, 2,394 eGenes and 108,782 *cis*-eQTLs were detected specifically in the NAFLD group (Fig. 1B-C). In comparison, 484 eGenes and 8,181 *cis*-eQTLs were detected as specific to the no-NAFLD group. Like the liver-eQTLs, NAFLD-eQTLs clustered throughout the whole genome (Fig. S3). Both eQTL sets showed high enrichment of eSNPs in gene bodies and proximity to the gene starts and ends (Fig. S4A). Relative to intergenic regions, we also detected enrichments in UTRs, introns, and ncRNAs, which are putative functional regions (Fig. S4B). We additionally performed a genome-wide functional enrichment analysis of liver-eQTL and NAFLD-eQTL sets using GREGOR[28] and observed that eSNPs were significantly enriched in known transcription factor binding and histone modification sites (Fig. S4C and S4D). Enrichment in these regulatory regions provides further evidence that the eSNPs are functionally relevant to gene expression and regulation in the liver. The numbers of tissues that expressed our eGenes were comparable to those for GTEx liver eGenes (Fig. S4E). And in the liver, our eGenes displayed higher expression than non-eGenes (*P* < 2.2 × 10^−16^, Welch’s *t*-test; Fig. S4F), suggesting a strong functional relevance. After additional filtering processes described in Fig. 1B and 1C, 242 NAFLD-eQTLs that alter expression in the NAFLD group but not in the no-NAFLD group, were selected for further analyses (Fig. 1B-1E and Table S4).

### AGXT2 expression is regulated by rs2291702 in NAFLD

Among the 242 NAFLD-eQTLs, we sought to identify loci that are biologically relevant and may contribute to NAFLD pathogenesis. We focused on the *AGXT2* locus as it is the second strongest signal after a lncRNA (*RP11-469A15.2)* and exclusively expressed in liver and kidney. One of the eSNPs, rs2291702, forms a significant *cis*-eQTL in the NAFLD group (*P* = 7.21 × 10^-9^), but not in no-NAFLD (*P* = 0.38) or GTEx liver (*P* = 0.31) sets (Fig. 2A). This significance persisted after adjusting for BMI and HOMA-IR in addition to age and sex (Fig. S5). Furthermore, rs2291702 also showed a clear association (*P* = 3.32 × 10^-5^) between its genotypes and *AGXT2* expression level in an additional Korean cohort (*n* = 162; Fig. 2B and Table S2). Therefore, combining this independent cohort with our subjects (*n* = 287) yielded stronger evidence that the eSNP functions in the NAFLD status (*P* = 3.69 × 10^-12^; Figure S6A and S6B). As a result, the difference in *AGTX2* expression between NAFLD and no-NAFLD (*P* = 3.47 × 10^-6^) was largely attributed to CC carriers (*P* = 4.79 × 10^-6^) and not to non-CC carriers (*P* = 0.16; Fig. 2C).

**Figure 2.**
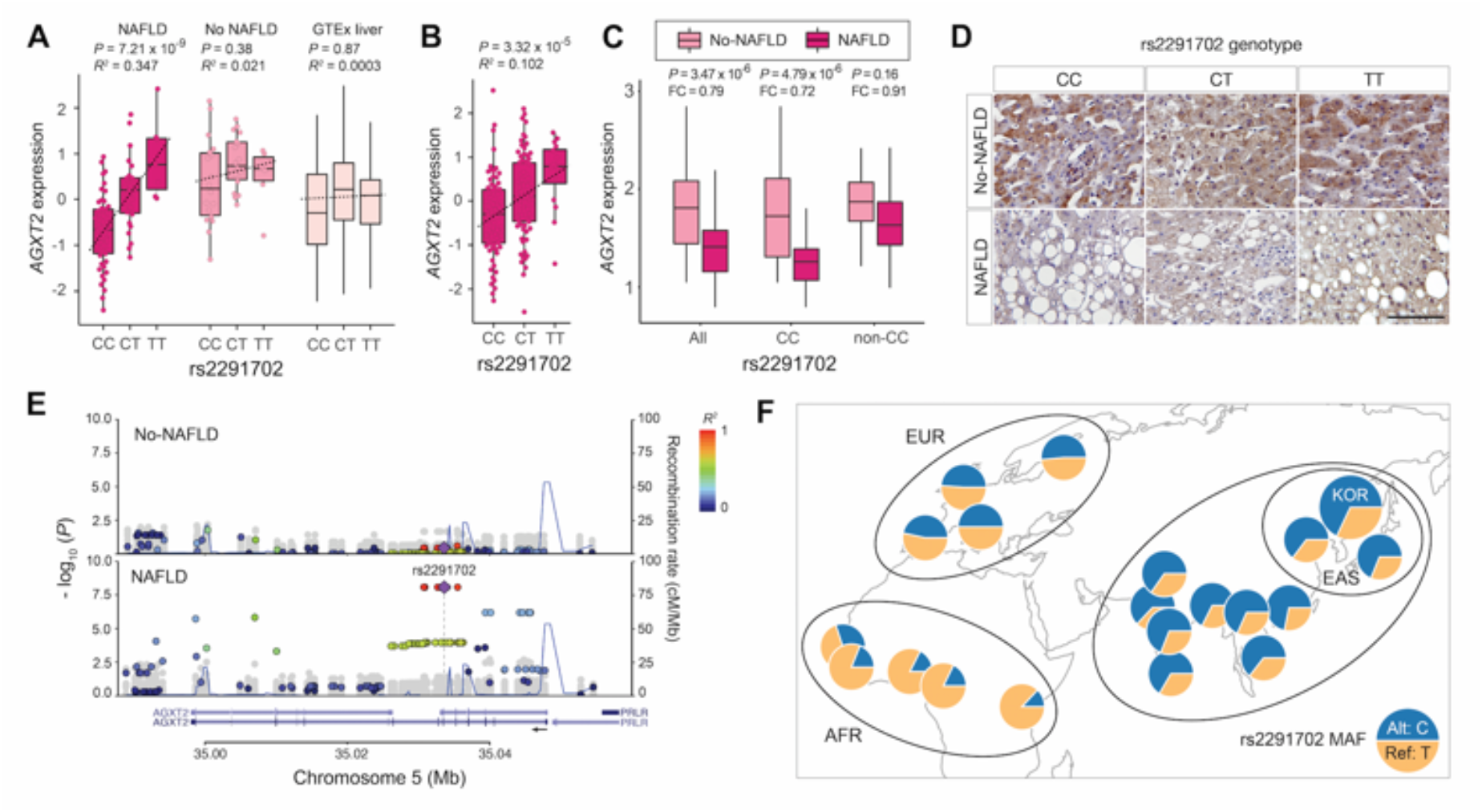
*AGXT2* expression is regulated by rs2291702 in a genotype-dependent manner. (A) *AGXT2* expression in NAFLD, no-NAFLD and GTEx liver samples. (B) *AGXT2* expression in the replication cohort. (C) Difference in expression of *AGXT2* by rs2291702 genotype. (D) Immunohistochemistry of liver biopsy samples corresponding to each genotype-phenotype category. Scale bar: 100 μm. (E) Regional plots of the eQTL association between each SNP’s genotype and *AGXT2* expression level. *AGXT2* locus displaying association from no-NAFLD (upper panel) and NAFLD (lower panel) groups. Color represents *R^2^* values between rs2291702 and nearby SNPs. SNPs in grey denote signals calculated using genes other than *AGXT2._* Blue lines denote recombination rate collected from 1000 Genomes Project (Phase 3). The arrow shows direction of *AGXT2* transcription. (F) World-wide allele frequency distribution of rs2291702. Data obtained from 1000 Genomes.

Immunohistochemistry of AGXT2 was performed in human liver biopsy samples and confirmed that its protein expression pattern is consistent with the gene expression pattern, which is dependent on the disease state and rs2291702 genotype (Fig. 2D and S7). The top eight *AGXT2*-eSNPs lie in a linkage disequilibrium (LD) block that spans ~4.7 kb in the 4–7th introns of the gene (Fig. 2E), and these SNPs show variable allele frequencies (AF) across different populations. In the 1000 Genomes database, the reference T allele of rs2291702 predominates over the alternative C allele in African populations (mean AF = 0.802), and has a frequency that is roughly half in European populations (mean AF = 0.511), whereas it is minor in East Asians and Koreans (AF = 0.327 and 0.330, respectively; Fig. 2F). This observation implies a potential population-specific role of these SNPs along with differential susceptibility to the *AGXT2*-dependent NAFLD pathway.

### Role of AGXT2 in NAFLD progression

*AGXT2* encodes a mitochondrial alanine-glyoxylate aminotransferase, which is responsible for systemic regulation of metabolites such as asymmetrical and symmetrical dimethylarginine (ADMA, SDMA), and BAIB.[19–21] It is enriched in the liver and kidney (Fig. 3A), and within the liver, it is mainly expressed in hepatocytes, as evidenced by single-cell RNA-seq analysis of the human liver (Fig. 3B)[29] and a western blot in HepG2 cells (Fig. 3C). In comparison, AGXT2 is expressed in a low amount in LX-2, a human hepatic stellate cell line (Fig. 3C). *AGXT2* expression is significantly correlated with pathological and clinical features such as the degree of steatosis, ballooning, fibrosis, and lobular inflammation, and the levels of hyaluronic acid (HA), alanine aminotransferase (ALT), aspartate aminotransferase (AST), and HOMA-IR (Fig. 3D, 3E and S8). As expected, this effect was more evident in rs2291702:CC carriers, reflected by higher *R^2^* values than those of non-CC carriers, whereas rs2291702:non-CC carriers displayed no significant correlations between the histological severity of NAFLD and *AGXT2* expression (Fig. 3E, Table S5 and S6). Therefore, it is plausible to postulate that *AGXT2*-eSNPs function as causative variants in NAFLD, and the alteration of *AGXT2* expression by the rs2291702 genotype modifies NAFLD pathogenesis.

**Figure 3.**
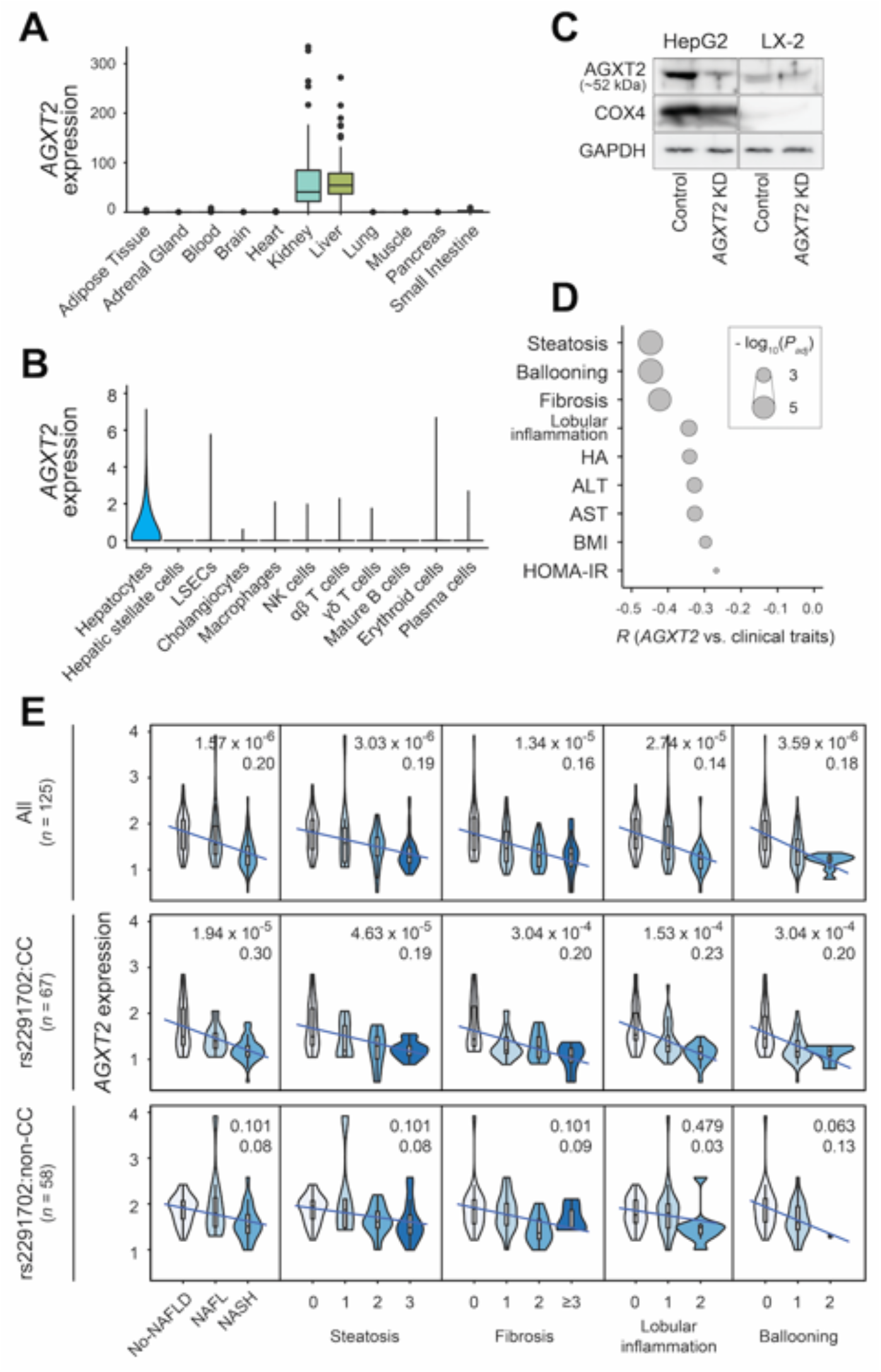
Hepatocyte-specific expression of *AGXT2* is associated with liver steatosis and fibrosis. (A) *AGXT2* expression in GTEx tissues. (B) *AGXT2* expression in human liver cell types analyzed at the single-cell level in the work of MacParland et al [29]. LSEC: liver sinusoidal endothelial cell. (C) AGXT2 expression in HepG2 and LX-2 cell lines. (D) Association of histological and metabolic parameters with *AGXT2* expression. (E) Correlation between the histological severity of NAFLD and *AGXT2* expression using all participants (top row), rs2291702:CC carriers (middle row) and rs2291702:non-CC carriers (bottom row). In each plot, the upper number indicates FDR-adjusted P-value and the lower number shows *R^2^* value.

Despite the growing evidence that AGXT2 and its substrates may play an important role in the pathogenesis of cardiovascular and metabolic syndromes,[17,18] the exact role of *AGXT2* in NAFLD is yet unclear. Therefore, we investigated whether altering expression of the gene will allow us to elucidate its function in regulating NAFLD progression. As the gene was downregulated in the NAFLD group (Fig. 2), we first tested whether the reduction of *AGXT2* may contribute to NAFLD development. *Agxt2* knockdown shRNA was injected into four nine-week-old mice on a normal diet. Seven days after the injection, histological analysis revealed that *Agxt2*-knockdown mouse livers featured an increase in collagen deposition (Fig. 4A and S9A). We also observed an increase in serum AST/ALT levels and hepatic transcript levels of fibrogenesis (*Col1a1* and *±SMA)*, inflammation (*Tnf±, Il-1²*and *Cd36)*, and adipogenesis (*Adrp* and *Acaca*), reflecting the NAFLD status (Fig. 4B and S10A-S10D).

**Figure 4.**
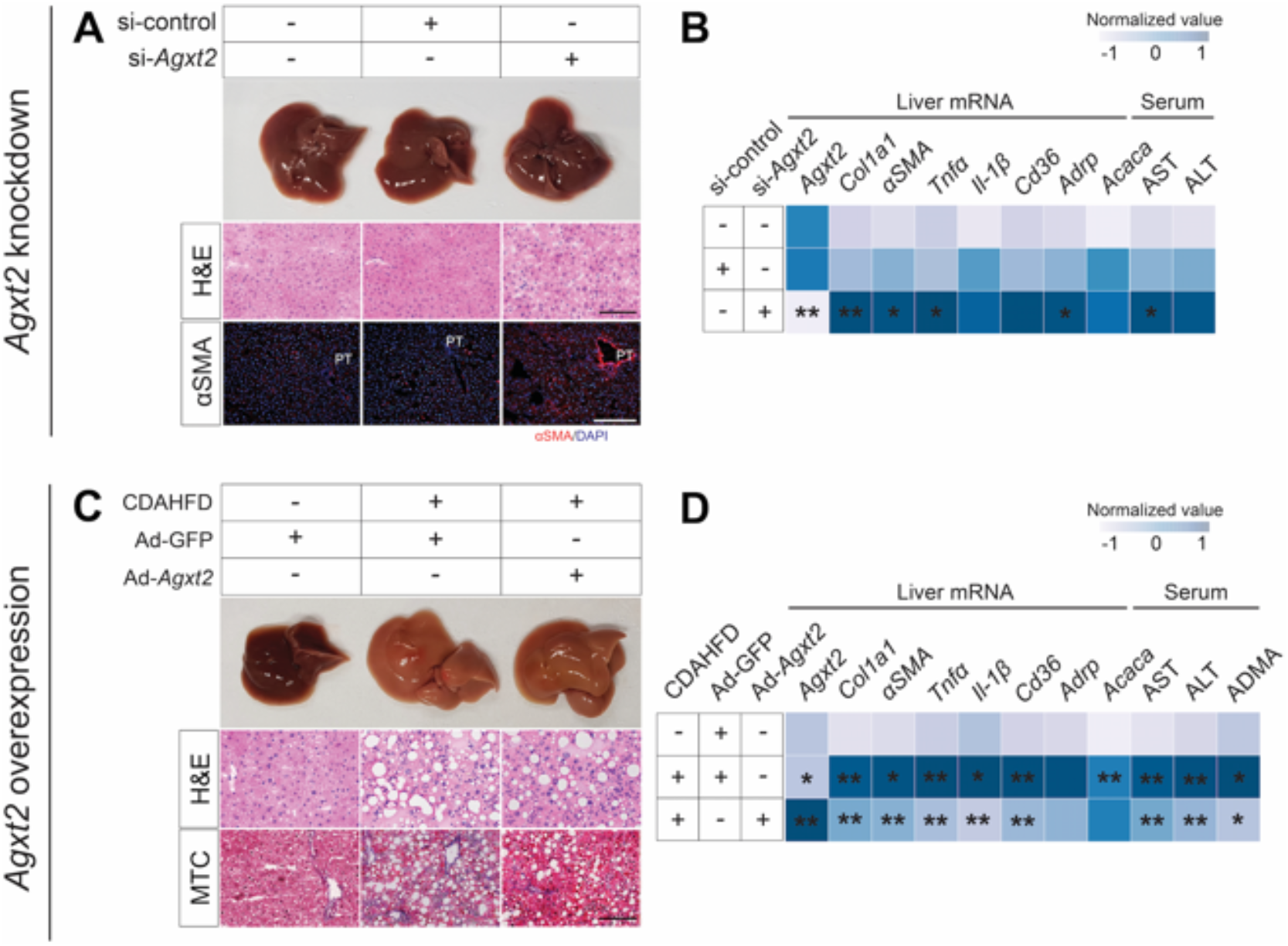
*Agxt2* knockdown aggravates NAFLD phenotypes and its overexpression ameliorates them. (A, C) Representative gross pictures, H&E, αSMA and MTC staining on sections of mouse livers. PT, portal triad. Scale bars: 200 μm. (B, D) Heatmaps representing normalized serum AST/ALT and ADMA levels and mRNA levels in *Agxt2* knockdown and overexpression mice. * *P* < 0.05, ** *P* < 0.01; independent *t*-test.

Next, we overexpressed *Agxt2* in four mice on CDAHFD, a widely-used NASH-fibrosis animal model.[30] Whereas the *Agxt2-* or *GFP*-injected mice were comparable in terms of fat accumulation (Figs. 4C), Masson’s trichrome (MTC) staining revealed the reduction of collagen deposition with the increased *Agxt2* dosage (Fig. 4C and S9B). The mice expressing *Agxt2* also displayed a decrease in serum AST/ALT levels and hepatic transcript levels of fibrogenesis, inflammation, and lipogenesis (Fig. 4D and S10E-S10H), indicating a protective role of AGXT2 in NAFLD progression. Moreover, *Agxt2*-overexpressed mice demonstrated a significant reduction in serum ADMA level, one of the main substrates of AGXT2

(Fig. 4D). Taken together, the knockdown and overexpression results suggest that *Agxt2* ameliorates fibrogenesis in the NAFLD mouse model, but show less pronounced effects on lipid accumulation and the degree of steatosis in the liver. In conclusion, AGXT2 protects from NAFLD progression by counterbalancing the fibrogenesis process.

### Alteration of the transcriptome in Agxt2 knockdown mice

Next, we investigated whether the reduction of *Agxt2* can genetically mimic human NAFLD features. We obtained liver samples from *Agxt2* knockdown (*n* = 2) and control (*n* = 2) mice, profiled transcriptional alteration by *Agxt2* depletion, and compared them with our human NAFLD transcriptomes (Fig. S11A). Of note, these mice were on a normal diet. We found a significant enrichment in the number of overlapped DEGs between human NAFLD patients and *Agxt2* knockdown mice (*P* = 3.78 × 10^-3^ for 163 concordantly regulated genes between human and mouse and *P* = 1.0 for 41 discordant genes; Monte-Carlo simulation), implying that the *Agxt2* knockdown mouse model harbors similar physiological features to human NAFLD status. Next, we performed gene ontology analyses to elucidate the functional basis of these DEGs. Both DEG sets exhibited NAFLD-related terms in common, but enrichments in the concordant genes more prominently featured metabolic functions (*e.g*., amino acid and lipid metabolic processes) (Fig. S11B). This supports that mouse transcriptomic changes resulting from the reduction of *Agxt2* resemble those observed in human NAFLD patients with metabolic abnormalities.

### AGXT2 knockdown induces ER stress and cell death

Based on the protective effects of Agxt2 against lipotoxicity of hepatocytes (Fig. 4 and S10), we sought to elucidate *in vitro* effects of *AGXT2* reduction (see supplementary method). Knockdown of *AGXT2* in the HepG2 cells (as demonstrated by western blot; Fig. 3C) sensitized cell death after palmitate treatment (*P* < 0.005, ANOVA with the Tukey’s test; Fig. 5A) and increased cellular levels of ER stress (GRP-78 and CHOP) and apoptosis markers (cleaved Caspase-3 and PARP) as detected by Western blot (Figs. 5B and S9C). Meanwhile, a reduced expression of *AGXT2* increased mitochondrial superoxide generation (*P* < 0.01, ANOVA with Tukey’s test; Fig. 5C), which was enhanced by the addition of palmitate. We also observed reduced mitochondrial integrity and oxygen consumption rate by the reduction of *AGXT2* (Fig. S12A-S12C). There were no differences in cell proliferation (Fig. S12D). Similar patterns were observed in murine primary hepatocytes, in which ER stress markers were increased by the addition of palmitate at 300 μM (Figs. 5D and S9D). The expression was somewhat reduced at 500 μM, reflecting our assumption that alternative pathways may be activated due to high PA level. As increased hepatic ER stress is known to cause cell death and fibrogenesis,[31] these results suggest that decreased AGXT2 exacerbates liver fibrosis by increasing ER stress-mediated hepatocyte death in NAFLD.

**Figure 5.**
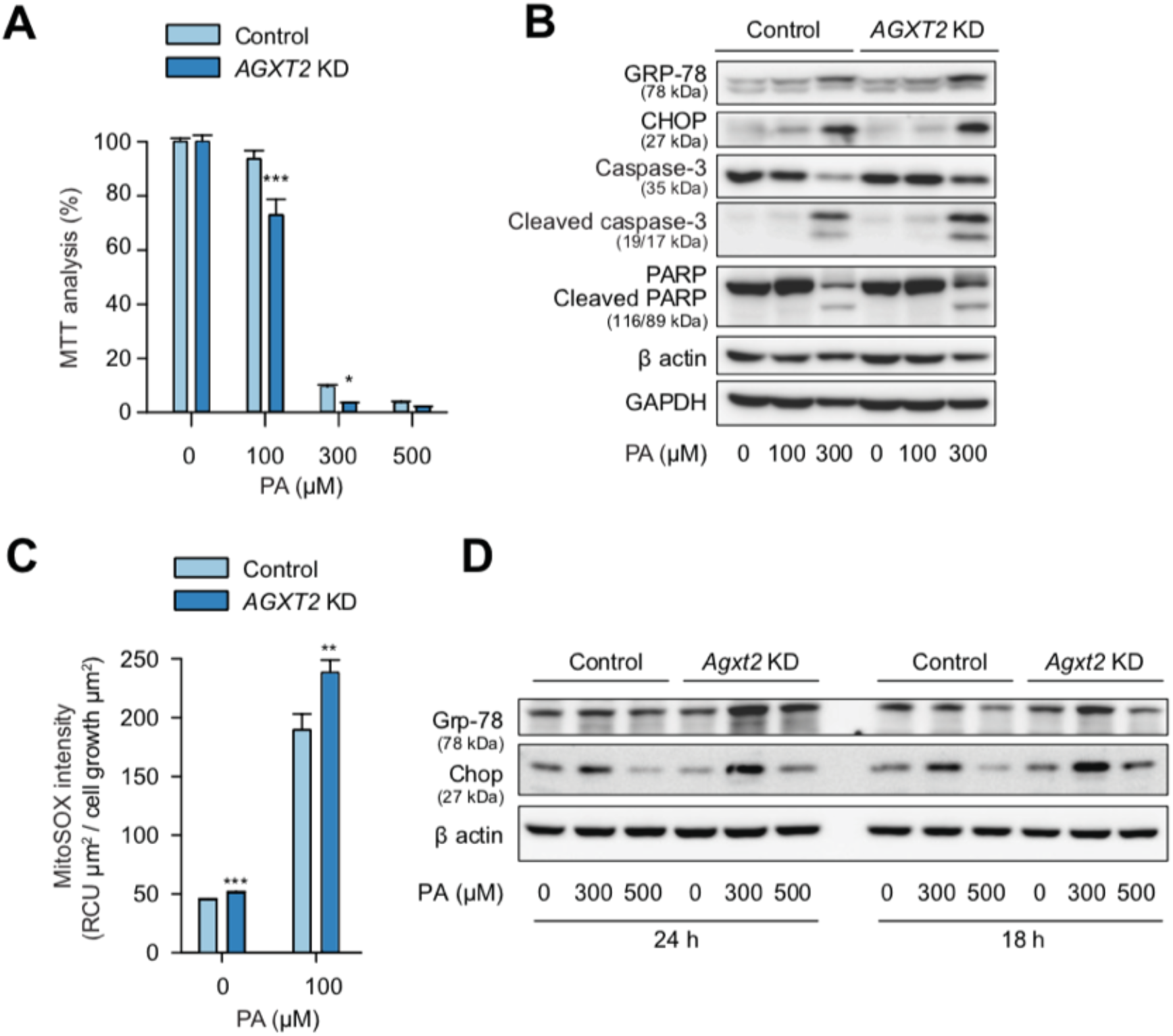
Downregulation of *AGXT2* induces ER stress and hepatocyte cell death under palmitate treatment. (A) Measurement of oxidative stress-mediated cell death by MTT analysis in HepG2 cells treated with PA for 48 hours (*n* = 4). (B) Western blot of ER stress markers in HepG2 cells by *AGXT2* knockdown and the addition of PA for 24 hours. (C) Measurement of mitochondrial ROS in HepG2 treated with PA (*n* = 3) for 24 hours. (D) Western blot of ER stress markers in mouse primary hepatocytes by *Agxt2* knockdown (24 hours) and the addition of PA (additional 18 or 24 hours). Values represented as mean ± SD, *\*P* < 0.05, \*\**P* < 0.01, \*\*\**P* < 0.005; ANOVA.

## Discussion

To accommodate the complex and polygenic nature of NAFLD, and to enhance the ability to identify genes underlying its pathomechanism, we present a disease-specific eQTL (*i.e*., NAFLD-eQTL) mapping pipeline. Unlike conventional eQTL approaches, our pipeline enabled us to identify eQTLs and associated genes that are active in the NAFLD environment. We confirmed that altered transcriptional activity depends on rs2291702, which is located in *AGXT2* and constitutes one of our top NAFLD-eQTLs. Using both mouse and cell models, we validated a NAFLD-preventive effect from *AGXT2*. Transcriptional changes of NAFLD-associated genes and pathways in *Agxt2*-knockdown mice on a normal diet indicate that suppression of *Agxt2* phenocopies human NAFLD with respect to gene expression. Lastly, at the cellular level, we present evidence that the reduction of *AGXT2* causes ER stress and hepatic cell death.

A number of studies have attempted to map healthy liver-eQTLs, revealing eQTLs that are active in normal physiological status.[22,32,33] Notably, a recent meta-analysis from 1,183 individuals detected that approximately 75% of all genes are associated with *cis*-eQTLs, consistent with a GTEx study.[27,34] In contrast with previous eQTL approaches using liver tissues,[32–36] our approach enables: (i) identification of eQTLs with disease risk, (ii) ranking eQTLs by effect size in the diseased state relative to the normal, and (iii) pinpointing individuals that harbor specific genotypes that are more or less susceptible to disease. For example, our analysis demonstrated that rs2291702:CC carriers display *AGXT2* downregulation and are more prone to the development and progression of NAFLD compared to others (*i.e*., CT/TT carriers). This feature does not necessarily selects SNPs that were also significant in GWAS, because we subdivided the cases and controls and this concomitantly reduces power. Nevertheless, this feature might be utilized in selecting proper target individual groups when developing and prescribing molecular targeted agents against NAFLD. A recent study demonstrated a similar pattern of the *MBOAT7* variant.[37] They observed that rs641738, a SNP near *MBOAT7* with significance from GWAS, confers the risk of liver fibrosis in a genotype-dependent manner.[37] This study and ours both highlight the functional implication of genetic variants that are active in a certain physiological condition. In addition, our approach illuminates the power of using non-European cohorts in discovering novel genetic players, as the susceptible allele predominates only in East Asian populations. It would be worthwhile to test whether the effect of genotype on *AGXT2* expression is valid in other ethnic cohorts.

We found that reduced *Agxt2* induces phenotypical and transcriptomic profiles similar to those of human NAFLD (Fig. 4), with livers of *Agxt2* knockdown mice displaying an increased expression of fibrogenesis, inflammation, and adipogenesis markers (Fig. 4B). Conversely, increasing *Agxt2* in high fat-fed mice reversed the marker expression changes (Fig. 4D), suggesting a protective effect of the gene on NAFLD. At the cellular level, we observed that reducing *AGXT2* induces an increase in ROS level and ER stress-mediated cell death upon metabolic stress, which is a well-known cause of liver fibrosis. This observation raises a question – how does reduced *AGXT2* contribute to ER stress and hepatic cell death? Investigations on the physiological role of AGTX2 have mainly focused on the cardiovascular system, as one of its main substrates, ADMA, is the most potent endogenous NOS inhibitor.

Moreover, a knockout mouse model displayed cardiovascular phenotypes.[18] In light of such findings, we tested if altered ADMA due to *AGXT2* dysregulation may affect NO level in the liver. However, *AGXT2* knockdown in HepG2 cells did not alter NO status, as assayed by 3NT western blotting (data not shown). Also ADMA itself failed to worsen the ER stress induced by PA treatment, implying other or additional substrates may be engaged in this process. Another mechanism of *AGXT2* pathogenesis is through altering amino acid metabolism, as AGXT2 utilizes various amino acid metabolites as substrates or products. Indeed, we observed that the levels of known substrates and amino acids that are closely linked with the AGXT2 enzymatic activity are altered by the reduction of *AGXT2* (Fig. S13) in HepG2 cells. Although the causal relationship of amino acid dysregulation with NAFLD remains to be clarified, this observation is consistent with previous studies[38,39] and offers a plausible explanation of hepatic damage mediated by the reduced *AGXT2*. Lastly, the precise role of hepatic stellate cells in this process also remains as an open question, although we have observed similar changes of fibrosis and inflammation marker expression upon *AGXT2* knockdown or overexpression in LX-2 cells (Fig. S14).

We observed that *AGXT2*-eSNPs are functional in NAFLD. Furthermore, we discovered that the SNPs possess transcriptional regulatory activity of varying degrees when placed on a luciferase reporter, and the effect can influence across the LD block that they are located (Fig. S15). Nevertheless, precise molecular basis of the *AGXT2*-eSNP function needs to be further studied.

Despite the stringent filtering steps carried out in this study, it would be necessary to increase the sample size to achieve sufficient statistical power, and to systematically validate the function of each eQTL under both normal and affected status. Although our set of NAFLD-eQTLs possesses many previously-associated NAFLD genes (Table S7), repeating our approach at the single-cell level would provide further insight into how cell type-specific NAFLD-eQTLs behave and confer the risk of NAFLD.

Here we presented a proof-of-concept method in which a disease-specific eQTL was selected and its function was validated toward the identification of therapeutic targets, specifically proposing *AGXT2* as a novel druggable target against NAFLD. Given that NAFLD is a polygenic trait and a substantial portion of the NAFLD population is nonobese and probably considered to harbor substantial genetic susceptibility, it is expected that additional loci will be mapped with biological validation. Therefore, as we increase sample size to boost statistical power and perform eQTL analysis on NAFLD and healthy individuals, it will become more feasible to find potent therapeutic targets and prospects for clinical interventions in an individual-specific manner. To date, therapeutic clinical trials have been unsuccessful largely because they failed to consider the genetic heterogeneity of NAFLD patients. In this respect, our approach has substantial advantages for the NAFLD drug development process through selective enrollment of a high-risk NAFLD population carrying risk variants.

## Authors Contributions

W.K. and M.C. conceived and designed the study; S.K.J., J.L. and W.K. contributed to sample acquisition; T.Y., Y.L. and M.C. contributed to sample processing, quality control and data generation; T.Y., O.J., S.R., W.K. and M.C. contributed to the analysis and interpretation of the data; H.J.K., H.Y.K., H.S., H.-H.K., S.J., W.-I.J., G.-S.H., K.W.K. and J.W.K. contributed to the functional analyses and interpretation; T.Y., W.K. and M.C. drafted the manuscript; all authors read and approved the manuscript.

## Data availability statement

The data that support the findings of this study are available from the corresponding authors, upon reasonable request. The RNA-seq data from mouse livers are available from the Korean Nucleotide Archive (https://kobic.re.kr/bps/kona; Accession ID: PRJKA200013).

## Abbreviations

eQTL: expression quantitative trait loci
AGXT2: alanine-glyoxylate aminotransferase
NAFLD: nonalcoholic fatty liver disease
ER: endoplasmic reticulum
NASH: nonalcoholic steatohepatitis
GWAS: genome-wide association studies
MBOAT7: membrane bound O-acyltransferase domain containing 7
PRS: polygenic risk score
ADMA: asymmetric dimethylarginine
BAIB: 3-amino-isobutyrate
DMGV: dimethylguanidino valeric acid
NOS: nitric oxide synthase
NAFL: nonalcoholic fatty liver
DEG: differentially expressed gene
MAF: minor allele frequency
HWE: Hardy-Weinberg equilibrium
CDAHFD: choline-deficient, L–amino acid-defined, high-fat diet
TSS: transcription start site
TES: transcription end site
FDR: false discovery rate
AF: allele frequency
ChIP-seq: chromatin immunoprecipitation sequencing
3’UTR: 3’untransrated region
SDMA: symmetrical dimethylarginine
ALT: alanine aminotransferase
AST: aspartate aminotransferase
HOMA-IR: homeostasis model assessment of insulin resistance
MTC: Masson’s trichrome
H&E: hematoxylin and eosin
ROS: reactive oxygen species
PA: palmitic acid
BMI: body mass index
SBP: systolic blood pressure
DBP: diastolic blood pressure
GGT: gamma-glutamyl transferase
Adipo-IR: adipose insulin resistance index

## Acknowledgments

We thank the participants of the study and Jana Kneissl for technical assistants. Silvia Sookoian, Tae Soo Kim, Yoon Jung Kim, Sangmoon Lee and Yongjin Yoo provided critical comments.

