## Supplementary materials for "Disease-specific eQTL screening reveals an anti-fibrotic effect of AGXT2 in nonalcoholic fatty liver disease"

### Supplementary methods

*Subjects*

The eligibility criteria of the Boramae NAFLD registry were (i) ≥18 years old, (ii) bright echogenic liver on ultrasound scanning (increased liver/kidney echogenicity and posterior attenuation), and (iii) unexplained elevated alanine aminotransferase (ALT) levels above the reference range within the past 6 months. The following exclusion criteria were adopted: (i) hepatitis B or C virus infection, (ii) autoimmune hepatitis, primary biliary cholangitis or primary sclerosing cholangitis, (iii) drug-induced liver injury or steatosis, (iv) Wilson′s disease or hemochromatosis, (v) excessive alcohol consumption (male >30 g/day, female >20 g/day), and (vi) diagnosis of malignancy within the past year. Control liver tissues were obtained from subjects who underwent liver biopsy in a pre-evaluation for donor liver transplantation or in a characterization of solid liver masses that were suspected to be hepatic adenoma or focal nodular hyperplasia based on radiological evaluation without any evidence of hepatic steatosis. All patients were informed of the study protocol and provided a written and signed consent. NAFLD was defined as the presence of ≥5% macrovesicular steatosis. NASH was diagnosed based on an overall pattern of histological hepatic injury consisting of macrovesicular steatosis, inflammation, and hepatocellular ballooning according to the criteria of Brunt et al.[1] NAFLD activity scoring and fibrosis staging were performed following the Kleiner classification, and study subjects were subsequently categorized into no-NAFLD, NAFL, and NASH (Table 1, S1 and S2).[2] In the subsequent analyses, we considered no-NAFLD as the control group, and both NAFL and NASH as the

NAFLD group. This study was conducted in accordance with the provision of the Declaration of Helsinki for the participation of human subjects in research and was approved by the institutional review board of Seoul Metropolitan Government Boramae Medical Center.

*RNA-seq and data processing*

Total RNA was isolated from liver biopsy samples using the conventional TRIzol (Invitrogen, Carlsbad, CA) RNA extraction protocol. Total RNA was quantified and assessed on a BioAnalyzer (Agilent Technologies, Inc., Santa Clara, CA). The mean RNA integrity number (RIN) value was 8.4 (4.5–9.5). cDNA library preparation was performed using TruSeq Stranded Total RNA Sample Prep kit (Illumina, Inc., San Diego, CA). The libraries were sequenced on a HiSeq2500 platform at Theragen Etex (Suwon, Korea) to generate paired-end 100 bp sequence reads. Read alignment/mapping and quantification were performed using TopHat[3] and HTseq[4] with the human genome (hg19/GRCh37) based on GENCODE v19. Differentially expressed genes (DEGs) were called using the DESeq2 packages[5] with correction for sample batches. Genes that were not expressed in at least one-third of subjects were excluded, and expression values were quantile-normalized.

*Genotyping and data processing*

Genomic DNA was isolated from blood samples using QIAamp DNA Blood Mini kit (QIAGEN Inc., Germantown, MD) according to the manufacturer's protocol. The mean 260/280 ratio of DNA samples was 1.82 (0.97–1.98), and the mean 260/230 ratio was 1.59 (0.30–3.68). Genotyping was performed with the Illumina Infinium OmniExpress-24 kit ($n$ = 81) or Omni2.5-8 kit ($n$ = 44) at the Yale Center for Genome Analysis (North

Haven, CT) following the manufacturer’s protocol. Raw genotype files were processed using Illumina GenomeStudio, and the mean call rate was 0.995. SNPs that are covered in both arrays were used.

The genotyped samples were processed with exclusion criteria of low call rate (< 0.95) and cryptic relationships using genome identity-by-descent (PI_HAT > 0.05) scores from PLINK (v1.9).[6] Genotype data were matched with RNA-seq data using VerifyBamID.[7] PLINK was used to remove markers with genotyping call rate < 0.97, minor allele frequency (MAF) < 0.1, and Hardy-Weinberg equilibrium $P_{HWE} < 1.0 \times 10^{-4}$. Then the remaining 427,128 markers were subjected to imputation using IMPUTE2.[8] with 1000 Genome Project phase 3 haplotypes as the reference. Inclusion criteria for the imputed genotype calls are: missing rate < 0.03, MAF > 0.1, and $P_{HWE} > 1.0 \times 10^{-4}$.

*Transcription start and end site (TSS and TES) enrichment and region-based annotation*

We obtained TSS and TES information from GENCODE human GRCh37.p13 reference genome (https://www.gencodegenes.org) and compared distance from eSNP to TSS and TES of corresponding eGene. To identify distribution of eSNPs throughout genome, we used region-based annotation using ANNOVAR.[9]

*Immunohistochemistry (IHC)*

Paraffin-embedded liver tissues were cut at 4 μm thickness using a microtome, placed on the slide, and stained using a Discovery XT automated IHC stainer (Ventana Medical Systems, Inc., Tucson, AZ). Slides were incubated with anti-AGXT2 primary antibody (1:500, ab231815, Abcam, Cambridge, UK) for 32 minutes at 37°C, and a secondary antibody OmniMap anti-rabbit HRP for 20 minutes at 37°C. Slides were incubated in DAB (3,3'-Diaminobenzidine)+ $H_2O_2$ substrate for 8 minutes at 37°C and counterstained with Hematoxylin and Bluing reagent at 37°C. Reaction buffer (pH 7.6 Tris buffer) was used as a washing solution. Slides were examined using a brightfield microscope. DAB intensity was quantified by Fiji with Color Deconvolution plugin.[10,11]

*AGXT2 overexpression and knockdown in HepG2 and LX-2 cell line*

HepG2 (#88065, Korean Cell Line Bank, Seoul, Korea) and LX-2 cells were plated at a density of $2 \times 10^5$ cells/well. The plasmid was packaged with Lipofectamine 2000 (Invitrogen), and cells were transfected with either a control or AGXT2-overexpressing vector according to the manufacturer's instructions. For siRNA treatments, the cells were plated in 35-mm-diameter dishes 18-24 hours at a density of $8 \times 10^4$ cells/dish before transfection. Cells were then treated with control or AGXT2-siRNA constructs at a concentration of 50 nM in Opti-MEM using RNAiMAX (Invitrogen), according to the manufacturer's protocol.

*Histological analysis of experimental animals*

The mouse liver tissues were fixed in 10% neutral-buffered formalin. Following fixation, the liver was trimmed, embedded in a cryo-block, sectioned, and stained with hematoxylin and eosin (H&E), Masson's Trichrome (MTC) or α-smooth muscle actin (αSMA) antibody (Sigma, A2547). To visualize αSMA, anti-mouse Alexa Fluor® 594-conjugated secondary antibody (Jackson ImmunoResearch, 115-585-003) was used. Fluorescence images were acquired using an Olympus BX51 microscope equipped with a CCD camera (Olympus, Tokyo, Japan) and computer-assisted image analysis with DP2-BSW.

*RNA preparation and RT-qPCR using the mouse liver tissues*

Total RNA was isolated from mouse livers using TRIzol reagent (Invitrogen) according to the manufacturer's instructions. For RT-PCR, cDNA was synthesized from 5 μg of total RNA using random hexamer primers and Superscript reverse transcriptase III (Invitrogen). RT-PCR was conducted using SYBR Green Master mix (Applied Biosystems, Foster city, CA) in a total volume of 20 μL. Transcripts were detected by RT-qPCR with a Step One instrument (Applied Biosystems). PCR reaction condition is as follow: initial denaturation at 94 ºC for 5 min, followed by 40 cycles of 94 ºC denaturation for 30 sec, 55 ºC annealing for 30 sec and 72 ºC extension for 20 sec. The reactions were terminated by final denaturation step at 94 ºC for 30 sec. All data were normalized to 18S rRNA or β-actin, and the FCs were obtained using the ΔΔ-Ct method. All reactions were performed in duplicates. Relative expression levels and standard deviations were calculated using the comparative method.

*ADMA enzyme-linked immunosorbent assay*

Serum ADMA level in mice was measured using a Mouse ELISA Kit (MBS048288, MyBioSource, Inc., San Diego, CA) according to the manufacturer's protocol.

*Mouse transcriptome analysis*

Liver tissues were obtained from *Agxt2* knockdown and control mice and used in transcriptome analysis. RNA extraction, library construction, sequencing, and RNA-seq data analysis procedures are as described above. The ToppGene Suite was used for gene list enrichment analysis.[12]

*Knocking down AGXT2 in HepG2 cells and primary hepatocytes*

GIPZ™ non-silencing and AGXT2-targeting lentiviral shRNA particles were purchased from Dharmacon™ (Lafayette, CO) and used per the manufacturer's instructions. The particles were added to HepG2 cells and after 48 hours of transduction, cells were selected by 4 μg/ml of puromycin. Negative control and Agxt2-targeting small interfering RNAs (siRNAs) were obtained from Bioneer (Daejeon, Korea). siRNAs were transfected into murine primary hepatocytes using Lipofector-EZ (AptaBio, Korea). For knockdown, primary hepatocytes were incubated for 24 hours in siRNA-containing transfection-optimized media.

*Cell culture and palmitic acid treatment*

HepG2 cells were cultured in low glucose DMEM supplemented with 10% FBS, 100 units/mL penicillin, and 100 μg/mL streptomycin at 37.5°C in humidified 5% CO2. Primary hepatocytes were isolated from 8-week-old C57B6/N male mice, and cells with viability above 90% were used for the experiments. Non-recirculating 2-step perfusion method was used for the isolation (Klaunig et al., 1981). Hepatocytes were plated on collagen-coated plate and after incubation for 4 hours, culture medium was changed for treatment of palmitic acid (PA; P0500, Sigma, St. Louise, MO). PA was dissolved into 100% ethanol, diluted in DMEM containing 1% (w/v) fatty acid-free BSA (107758350001, Roche) to a final concentration and incubated for 30 minutes in 37°C before treated to cells.

*Measurement of cell viability and mitochondria isolation*

MTT (M2128, Sigma) assay was used to measure cell viability. Briefly, MTT (1 mg/mL dissolved in PBS) was diluted to a final concentration of 0.3 mg/mL with serum-free media. The cells in 24-well plates were incubated with the MTT containing media, away from light. After 2 hours of incubation, the media were removed by careful pipetting, and the resident cells were dissolved in 250 μL DMSO. The absorbance was measured at 590 nm with a TriStar microplate reader (Berthold Technologies, Bad Wildbad, Germany). Mitochondria were isolated from HepG2 cells using a reagent-based method (89874, Thermo Scientific).

*Western blot*

Western blotting was performed on HepG2 cells and primary hepatocytes. Proteins were transferred to nitrocellulose membranes and blocked with 5% skim milk for 1 hour at RT followed by incubation with each primary antibody. Then, membranes were washed and incubated with horse radish peroxidase–conjugated secondary antibody. The immune complexes were detected by using the Immobilon Western Chemiluminescent HRP Substrate (WBKLS0500, Merck Millipore, Billerica, MA). Protein levels were quantified by densitometry using software, Multi Gauge, V3.0 (Fujifilm, Tokyo, Japan).

*Fluorescence detection of mitochondrial reactive oxygen species (ROS)*

To investigate mitochondrial ROS, HepG2 cells were incubated in 5 μM MitoSox (M36008, Invitrogen) containing Hank's balanced salt solution (14025-092, Gibco) for 30 minutes at 37°C, away from light. MitoSox positive cells were detected and analyzed using Incucyte®. Intensity and area of red fluorescence were divided by phase area of each well.

*JC-1 assay*

Mitochondrial membrane-permeant fluorescent dye JC-1 (Thermo Scientific, Madison, WI) was used to detect changes in mitochondrial membrane potential, following the manufacturer's instruction. HepG2 cells were incubated in PBS containing 2 mg/ml JC-1 dye. After 30 minutes, cells were washed twice with PBS and analyzed using Incucyte®. Monomer and the J-aggregates were detected separately in green and red channels, respectively. The red to green emission ratio provided an estimate of mitochondrial membrane potential. Total integrated intensity (GCU or RCU × $\mu m^2$) of each channel was used to determine the red to green emission ratio.

*Measurement of mitochondrial respiration*

Oxygen consumption rate (OCR) was measured as an indicator of mitochondrial respiration by using Seahorse XFp Analyzer (Agilent). HepG2 cells were trypsinized and seeded into XFp cell culture mini plates (103022-100, Agilent) at 10,000 cells per well two days before the assay. Three technical replicates were used for each group. XF Assay Medium (102353-100, Agilent) was supplied with 5.5 mM glucose, 1 mM sodium pyruvate (11360-070, Gibco), and 2 mM L-glutamine (25030149, Gibco).

*RNA Preparation and RT-qPCR in human cell line*

Total RNA from primary hepatocytes was isolated using TRIzol® reagent (15596018, Thermo Fisher Scientific), and Oligo (dT) primers and reverse transcriptase (25081, iNtRON Biotechnology) were used to synthesize cDNA. The expression of AGXT2 were measured by RT-qPCR using a SYBR Select Master Mix (1708882, Bio-Rad) and Bio-Rad CFX Manager™ Software (Bio-Rad). 18S rRNA was employed as a reference for normalizing AGXT2 mRNA level.

*Liquid chromatography/mass spectrometry (LC/MS) analysis of amino acids*

Cells were quickly washed with cold PBS and then scraped into 2 mL of methanol/water mixture (8:2, v/v). Samples were vortexed vigorously for 1 minute and spun down at 17,500 g for 10 minutes. The supernatants were evaporated under a stream of nitrogen and reconstituted using 10% sulfosalicylic acid. The solution was derivatized with AccQ Tag 3X kit as following the manufacturer's protocol. For the ultra-performance liquid chromatography/triple-quadrupole mass spectrometry (UPLC/TQ-MS) analysis was performed using Acquity UPLC I-class systems equipped with Xevo TQ-XS mass spectrometer and electrospray ionization (ESI) source (Waters Corporation, Milford, MA). The TargetLynx Application Manager (Ver 4.2, Waters) software was used for data acquisition and analysis. LC separation was carried out using CORTECS UPLC T3 column (2.7 μm, 2.1 mm × 100 mm, Waters). The binary gradient system comprised 0.1% formic acid in water (solvent A) and 0.1% formic acid in acetonitrile (solvent B). The linear gradient used for elution and equilibrating the initial gradient for subsequent runs were as follows: 1% B from 0–0.67 minutes, 13% B to 1.33 minutes, 15% B to 3.67 minutes, 95% B to 4.33-5 minutes, 1% B from 5.1–7 minutes. Column temperature and flow rate were 45°C and 0.6 mL/minute, respectively. The auto-sampler temperature was maintained at 10°C, and an injection volume of the sample was 5μL. Samples were analyzed in single reaction monitoring (SRM) mode. The SRM transitions were performed with the following operational parameters: capillary voltage, 2kV; cone voltage, 20V; source temperature, 150°C; desolvation temperature, 650°C; cone gas flow, 150

L/hour; desolvation gas flow, 900 L/hour; collision gas flow, 0.15mL/minute; nebulizer gas flow, 7 bars. The intensity of whole cell amino acids was normalized to the total number of cells per group.

*Utilization of ENCODE data*

For epigenetic analysis of the *AGXT2* locus, we downloaded data tracks from the ENCODE portal (https://www.encodeproject.org/) with the following identifiers: CTCF ChIP-seq of HepG2 (ENCSR000BIE) over control (ENCFF804LNZ), CTCF ChIP-seq of hepatocyte (ENCSR252QYR) over control (ENCFF911ZCT), RAD21 ChIP-seq of HepG2 (ENCSR000BLS) over control (ENCFF941EXK), RAD21 ChIP-seq of liver (ENCSR230ZWH) over control (ENCFF110WCE).

*Plasmid Construction and Luciferase Assay*

To construct luciferase reporter plasmids with putative enhancers, we amplified patients' genomic DNA for each allele by PCR and prepared inserts with extension primers (Table S6). PCR products were inserted into SalI and BamHI double-digested pGL3-Promoter vector, using an overlap cloner (Elpisbiotech Inc., Daejeon, Korea). Resulting constructs were confirmed by Sanger sequencing. The dual luciferase assay was conducted using the Dual-Luciferase Reporter Assay System (E1960, Promega, Madison, WI). Briefly, HepG2 and LX-2 cells were transfected with the firefly-luciferase and Renilla-luciferase reporter plasmids for 24 hours, followed by oleic acid (OLA)/palmitic acid (PA) (500 μM and 250 μM each, bound with BSA) and TGF-β (2.5 ng/mL, Research and Diagnostic Systems, Inc., Minneapolis, MN) treatment. After 24 hours, samples were dissolved with 1x passive lysis buffer for 15 minutes, 20 μL of supernatant was transferred to a 96-well plate,

100 μL of luciferase assay reagent II was added, and firefly luciferase activity was measured. Then, 100 μL of Stop and Glo reagent was added and Renilla luciferase activity was measured. The ratio of firefly and Renilla signals was used to normalize transfection variation.

*Statistical analysis*

Data are presented as mean ± standard error of the mean or standard deviation and remarked in each figure. To evaluate the statistical significance of differences among no-NAFLD, NAFL, and NASH, the independent t-test, Mann–Whitney U-test, analysis of variance (ANOVA), or Kruskal–Wallis test was used for continuous variables and the chi-squared test was used for categorical variables. Statistical values from eQTL analysis were calculated based on linear regression under additive model which is implemented in MatrixEQTL package in R. Age and sex were used as covariates and for validation, BMI and HOMA-IR were additionally used if needed. All statistical significance of experimental results was evaluated with the independent t-test, Mann–Whitney U-test, and ANOVA. Significance was defined as $P < 0.05$. Multiple testing correction was done by FDR. All statistical analyses were conducted using R software version 3.5.1.

### Supplementary Figures

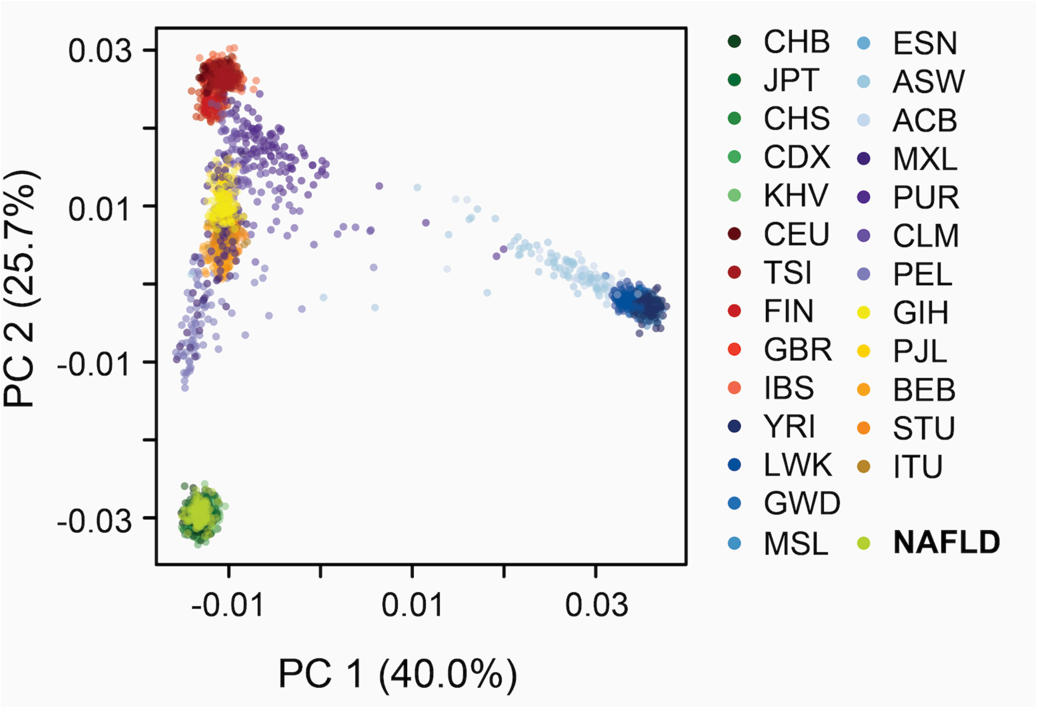

**Supplementary Fig. 1. Principle component analysis plot of our discovery cohort individuals (light green), plotted with the 1000 Genomes individuals.**

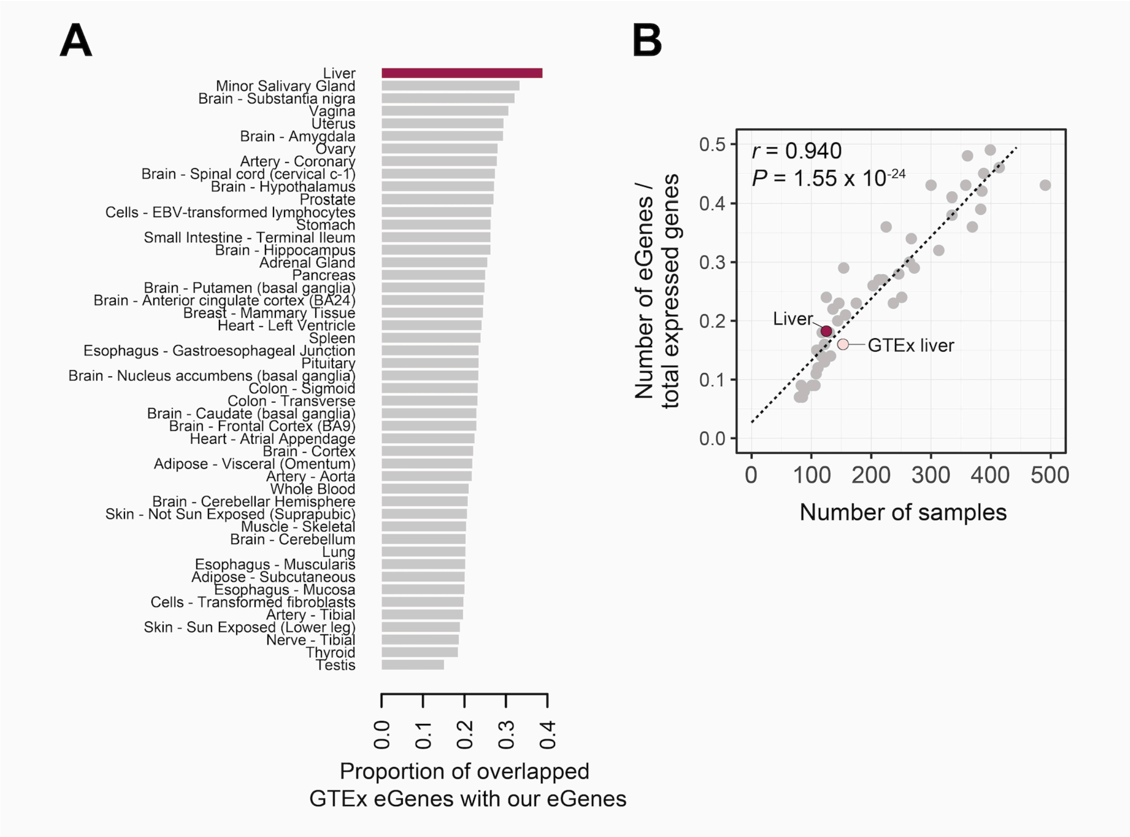

**Supplementary Fig. 2. Comparison of our NAFLD eQTLs with GTEx liver eQTLs.** (A) Proportion of eGenes from each GTEx set that overlapped with our liver cis-eQTL set. (B) Number of detected significant eGenes by sample size. "Liver" indicates our study.

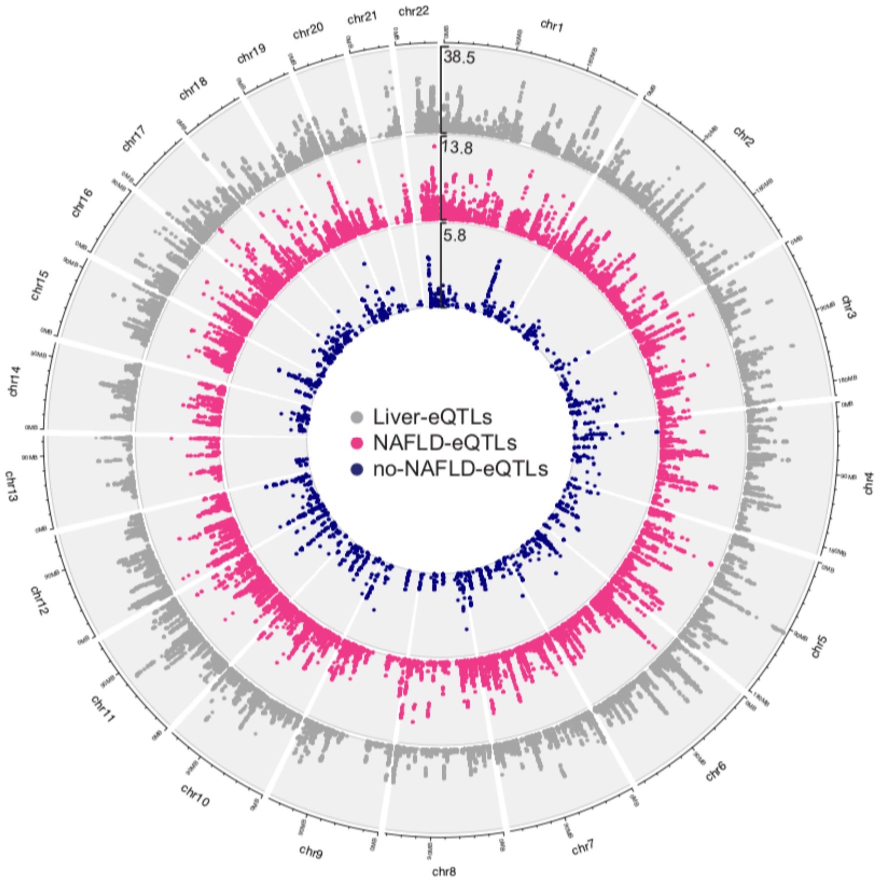

**Supplementary Fig. 3. Genome-wide eQTL distribution.** X- and Y-axis represent genomic position of eQTLs and -$\log_{10}(P_{adj})$, respectively. Visualized with shinyCircos.[13]

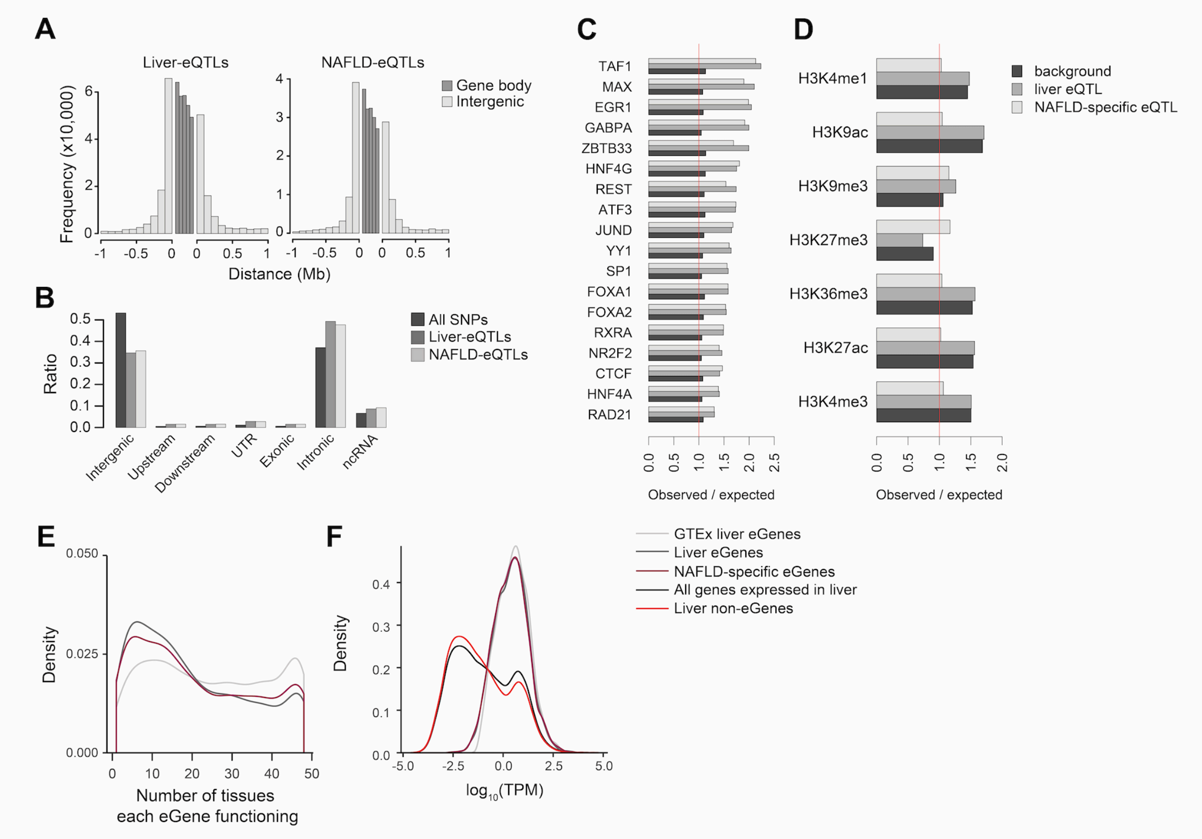

**Supplementary Fig. 4. Features of *cis*-eQTLs.** (A) Distribution of liver-eQTLs (left) and NAFLD-eQTLs (right) relative to TSS and TES of eGenes. Gene body region was divided into five relative bins. (B) Distribution of liver-eQTLs (darkgrey) and NAFLD-eQTLs (grey) by genomic annotation. (C) Overlap of liver-eQTLs and NAFLD-eQTLs by transcription factor binding elements, displayed by observed vs expected events. (D) Overlap of liver-eQTLs and NAFLD-eQTLs by histone modification signatures, displayed by observed vs expected events. (E) Distribution of the number of tissues where each eGene is expressed. (F) Distribution of expression levels in livers.

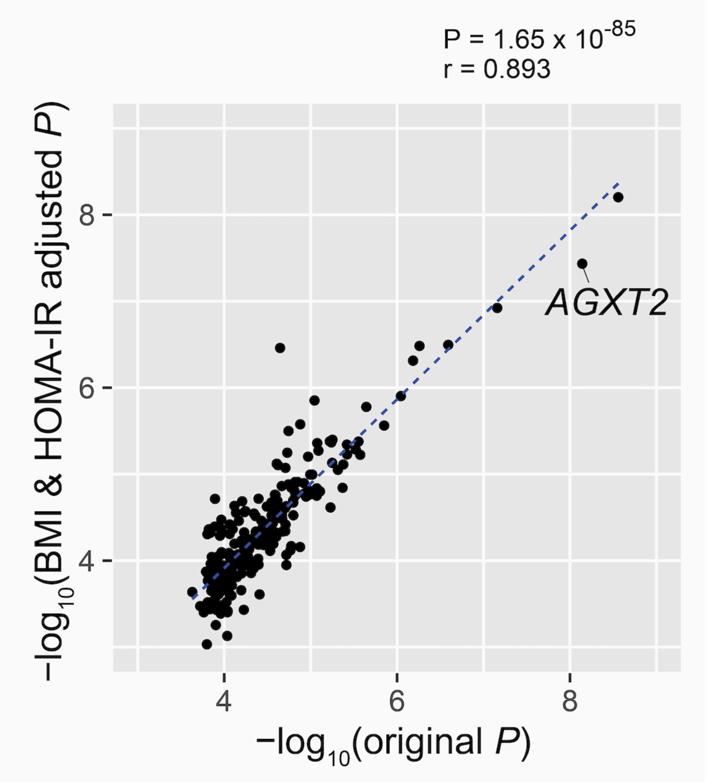

**Supplementary Fig. 5. Scatter plot of the 243 NAFLD-eQTLs comparing the *P*-values using age and sex as covariates (X-axis) vs age, sex, BMI and HOMA-IR as covariates (Y-axis).**

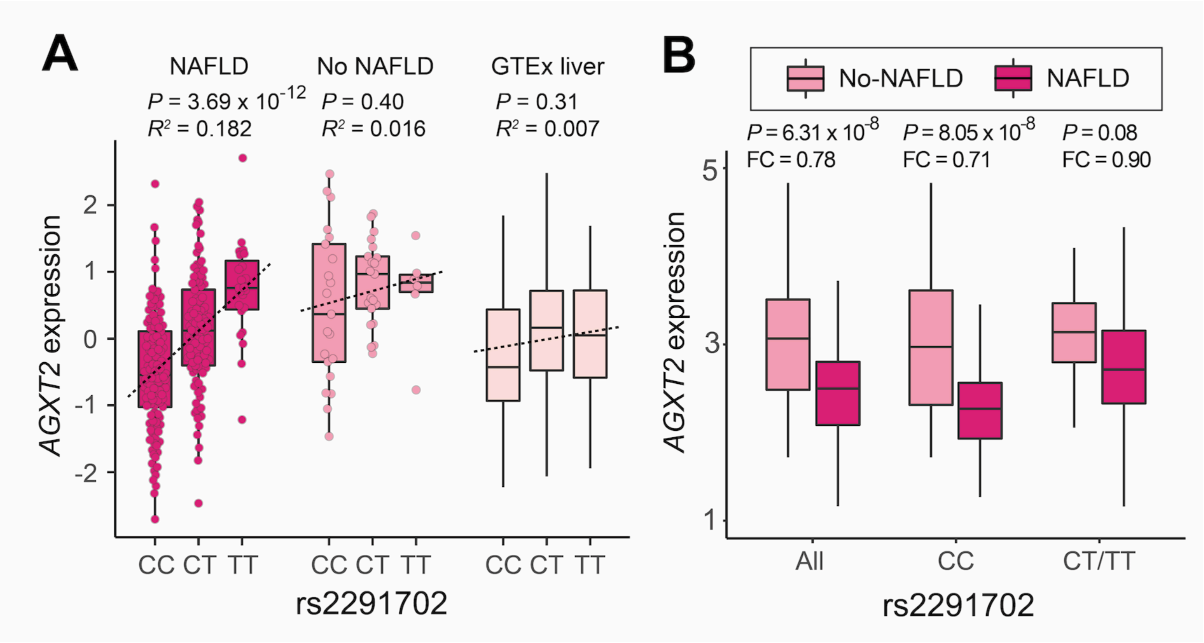

**Supplementary Fig. 6. *AGXT2*-eQTL in the combined set of 293 individuals.** (A) AGXT2 expression by rs2291702 genotype. (B) AGXT2 expression differences divided by rs2291702 genotype.

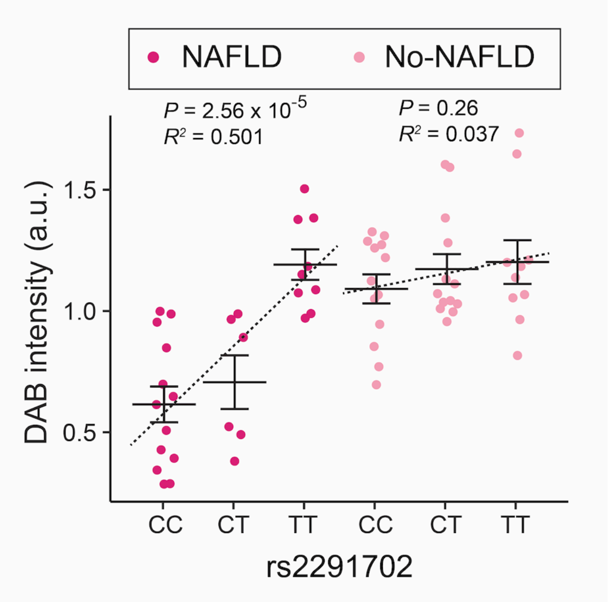

**Supplementary Fig. 7. DAB intensity quantified from AGXT2 immunohistochemistry images upon different genotypes and disease conditions.** a.u.: arbitrary unit.

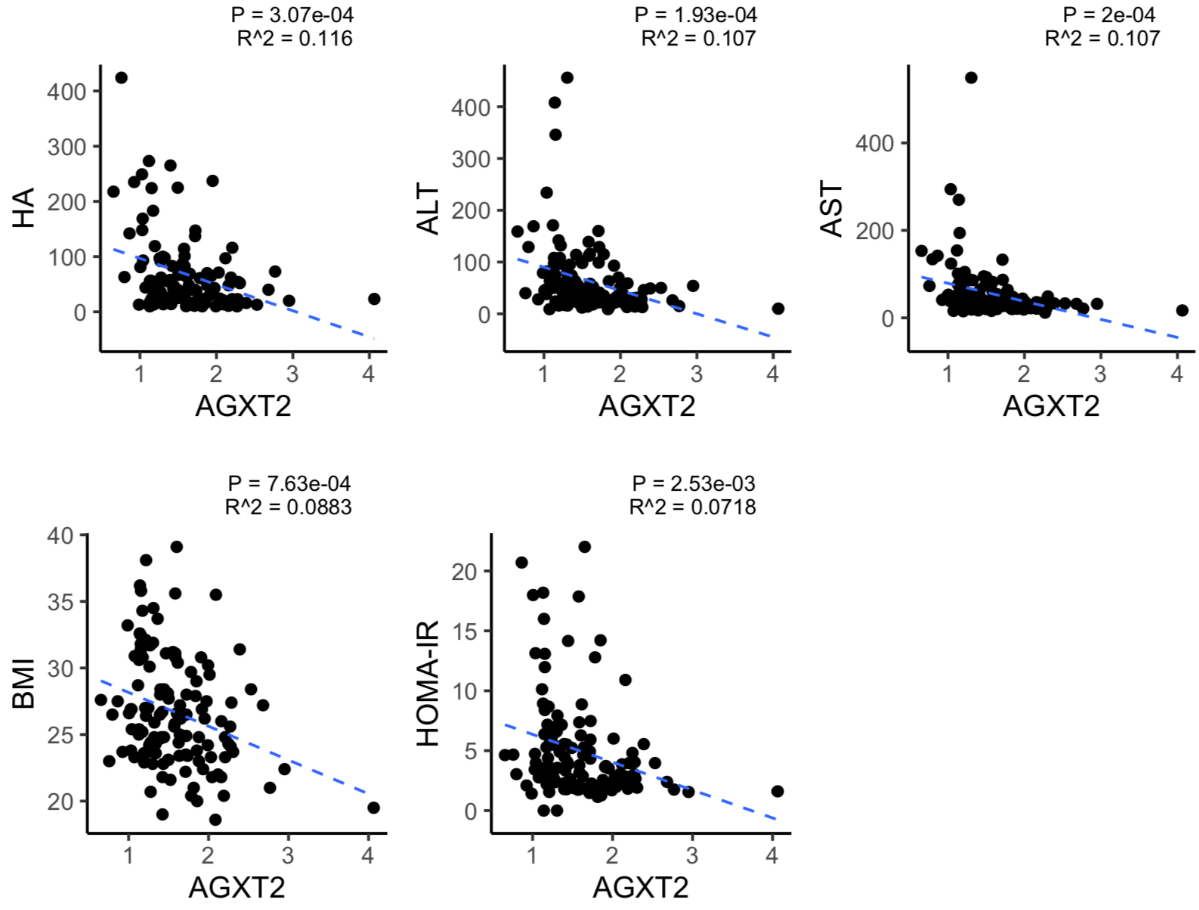

**Supplementary Fig. 8. Correlations between AGXT2 expression and clinical parameters in the NAFLD cohort shown in Figure 3D.**

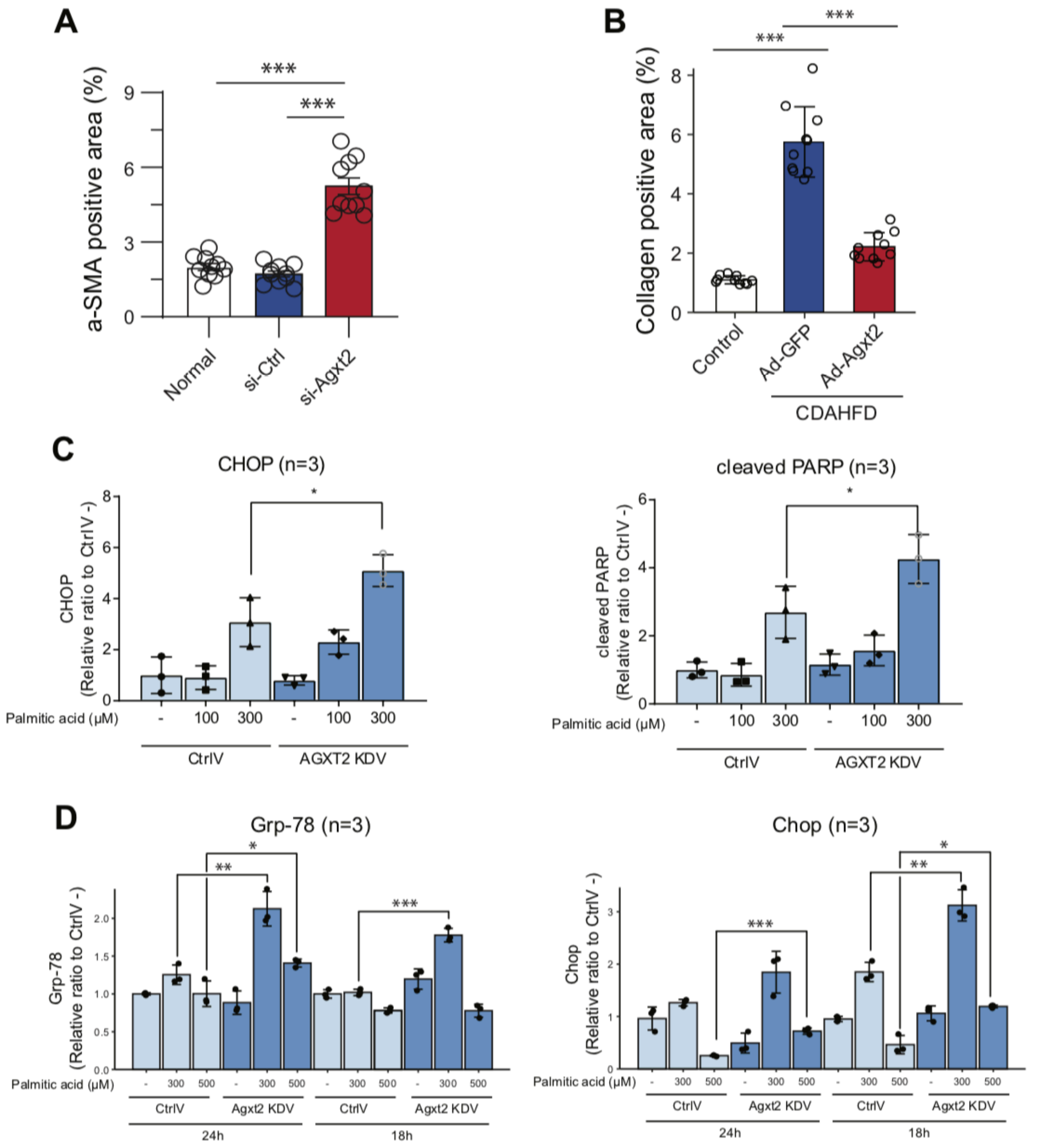

**Supplementary Fig. 9. Quantifications of results in Figs. 4A, 4C, 5B, and 5D.** (A-B) Quantification of αSMA signals in Fig. 4A (A) and collagen fiber in Fig. 4C (B). The stain-positive areas were analyzed in ten randomly chosen area per liver section. (C-D) Replication and quantification of the Western blot results shown in Figs. 5B (C) and 5D (D). ImageJ software was used for the quantification analyses. *. *P* < 0.05; **. *P* < 0.01; ***. *P* < 0.005.

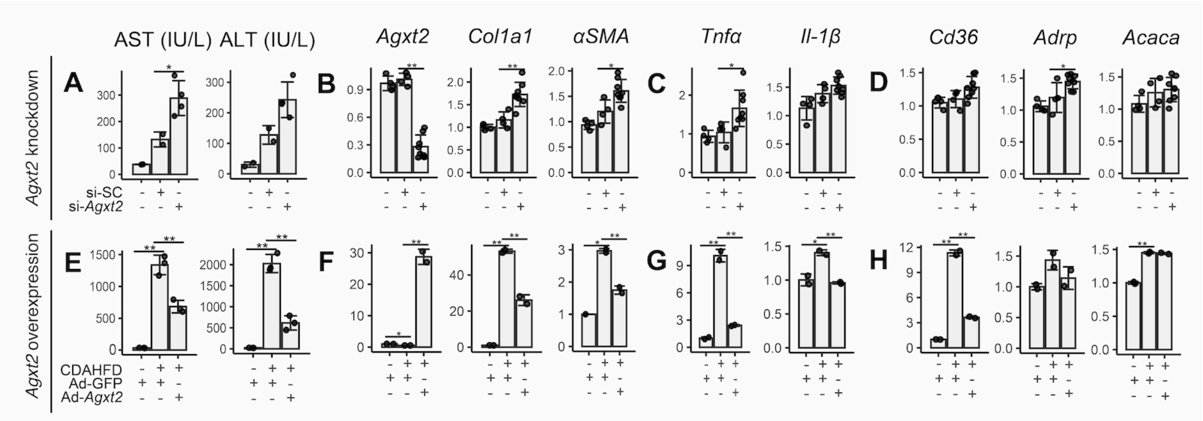

**Supplementary Fig. 10. Changes in liver injury markers and fibrogenesis, inflammation, and adipogenesis expressions by knockdown or overexpression of *Agxt2* in mice, as displayed as heatmaps in Figures 4B and 4D.** (A,E) AST, ALT levels in mouse serum. Relative expression of fibrogenesis (B,F), inflammation (C,G), and adipogenesis markers (D,H) in mouse liver. Upper and lower panels are from *Agxt2* knockdown and overexpression experiments, respectively. * $P < 0.05$, ** $P < 0.01$; independent t-test.

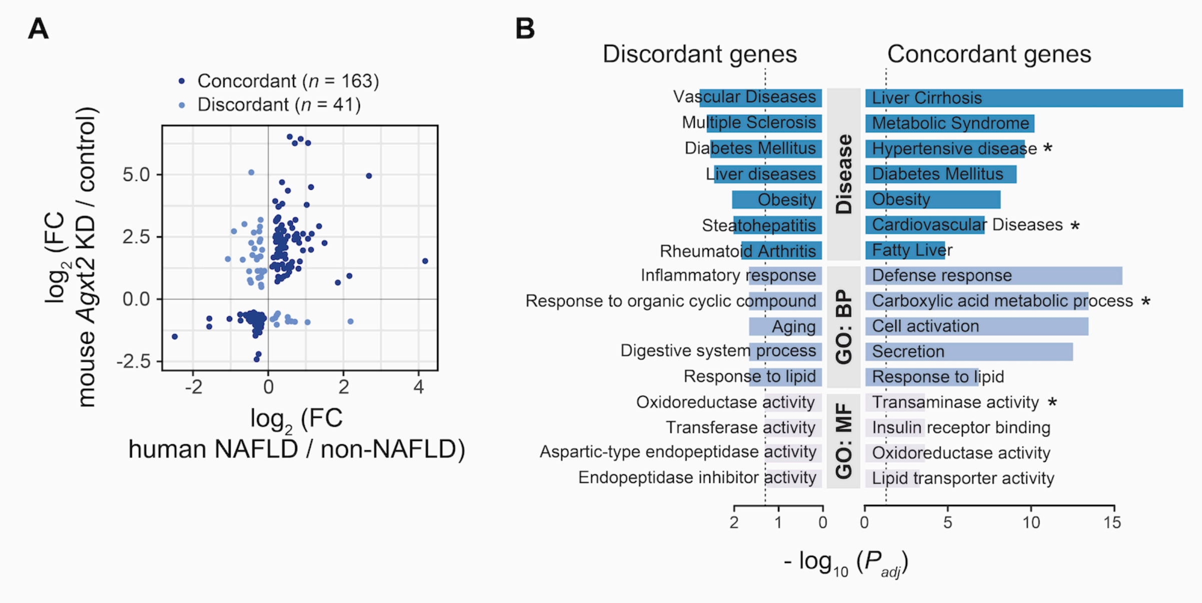

**Supplementary Fig. 11.** (A) Scatterplot of DEGs from mouse *Agxt2* knockdown and human NAFLD livers. (B) Gene enrichment analysis of concordant (right) and discordant (left) gene groups derived from (A). Terms with an asterisk are *AGXT2*-containing. Note the different *P*-value scales between the two groups.

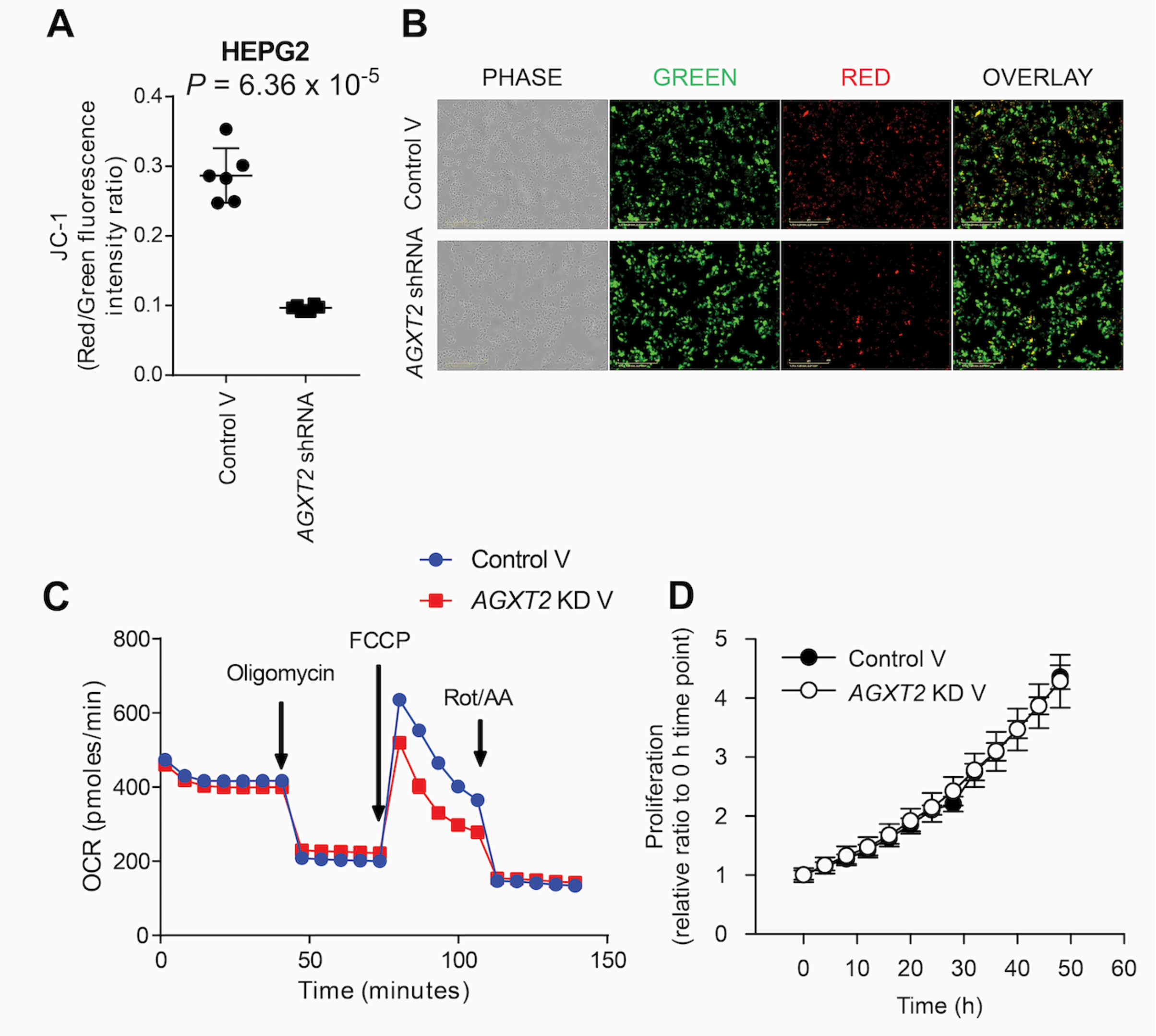

**Supplementary Fig. 12. Effects of AGXT2 knockdown on mitochondria, oxygen consumption, and cell proliferation.** Measurement of mitochondria polarity using JC-1 (A-B), oxygen consumption (C) and cell proliferation (D) in HepG2 cells.

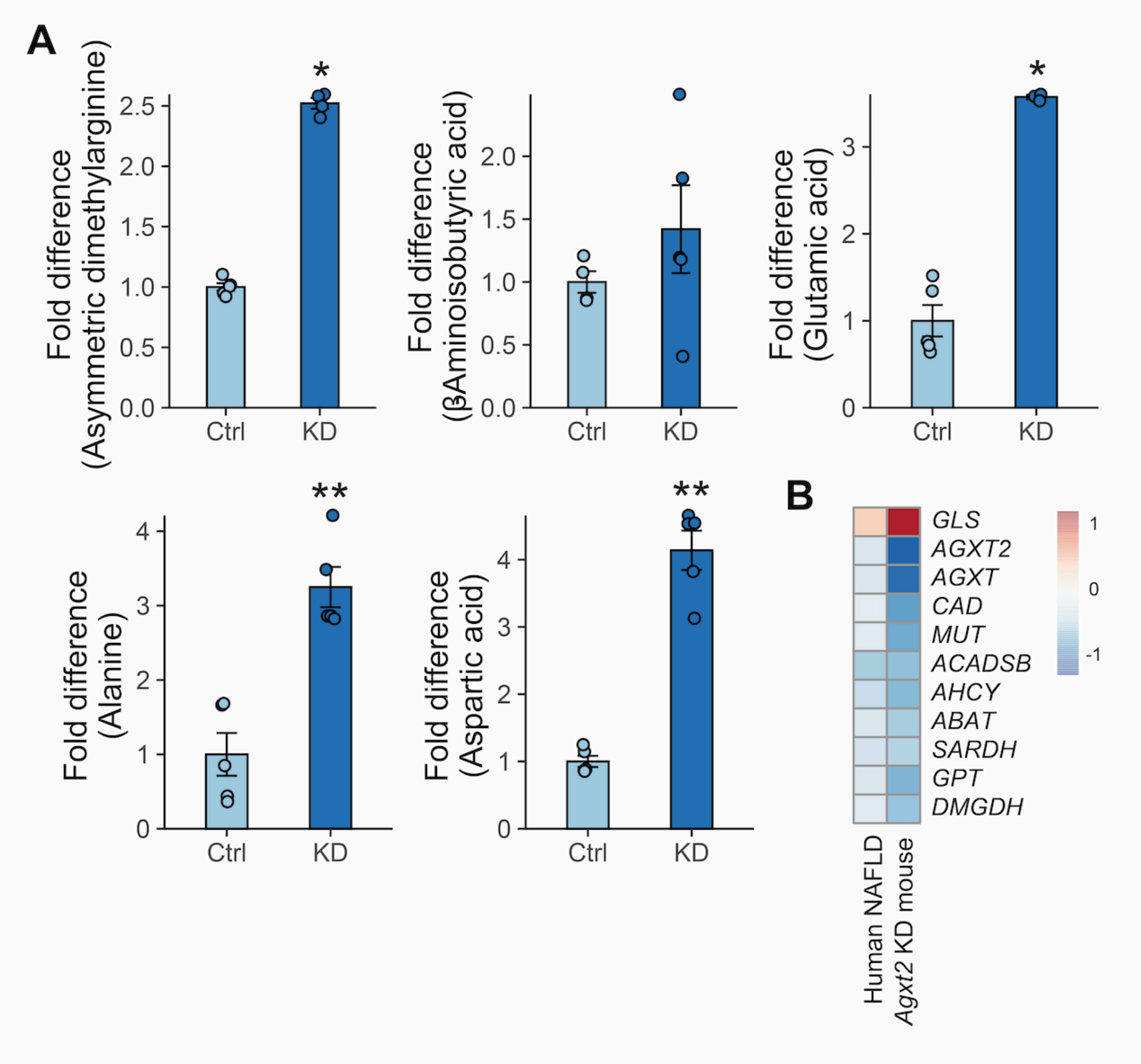

**Supplementary Fig. 13. Effects of AGXT2 knockdown on metabolic changes.** (A) Fold differences in AGXT2 substrate molecules and amino acids that are engaged in AGXT2 enzymatic activity between control and *AGXT2* knockdown HepG2 cells. (B) Genes involved in *AGXT2*-related amino acid metabolism pathways. Color represents expression fold change between no-NAFLD and NAFLD in human and *Agxt2* control and *Agxt2* KD in mouse. *$P < 0.05$, **$P < 0.01$; Mann-Whitney *U*-test.

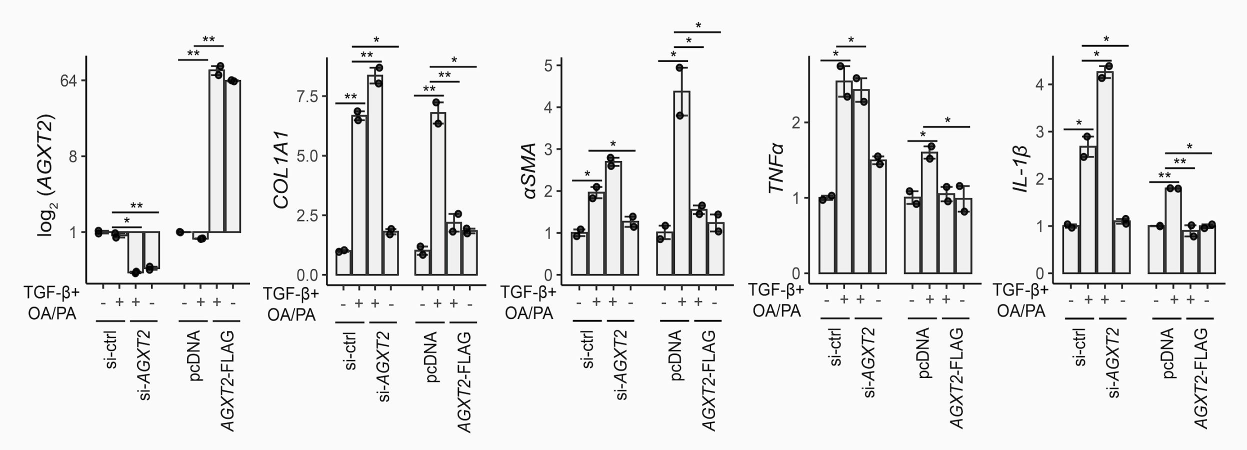

**Supplementary Fig. 14. Changes in profibrogenic and inflammatory markers by knockdown or overexpression of *AGXT2* in human hepatic stellate cell line LX-2.** Relative mRNA expression of fibrogenesis (*COL1A1 and αSMA*), and inflammation markers (*TNFα* and *IL-1β*) in LX-2 cell line. * $P < 0.05$, ** $P < 0.01$; independent *t*-test.

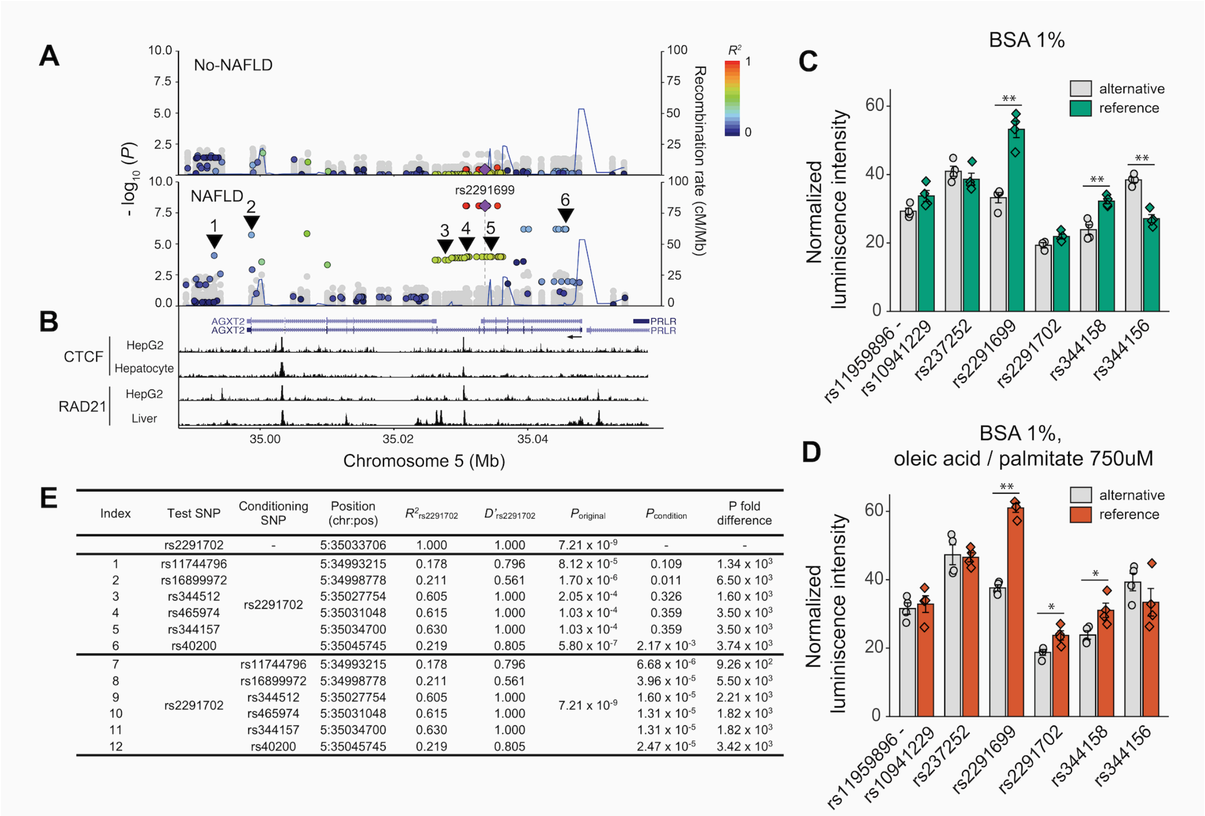

| Index | Test SNP | Conditioning SNP | Position (chr:pos) | $R^2_{rs2291702}$ | $D'_{rs2291702}$ | $P_{original}$ | $P_{condition}$ | P fold difference |
|---|---|---|---|---|---|---|---|---|
| | rs2291702 | - | 5:35033706 | 1.000 | 1.000 | $7.21 \times 10^{-9}$ | - | - |
| 1 | rs11744796 | rs2291702 | 5:34993215 | 0.178 | 0.796 | $8.12 \times 10^{-5}$ | 0.109 | $1.34 \times 10^{3}$ |
| 2 | rs16899972 | | 5:34998778 | 0.211 | 0.561 | $1.70 \times 10^{-6}$ | 0.011 | $6.50 \times 10^{3}$ |
| 3 | rs344512 | | 5:35027754 | 0.605 | 1.000 | $2.05 \times 10^{-4}$ | 0.326 | $1.60 \times 10^{3}$ |
| 4 | rs465974 | | 5:35031048 | 0.615 | 1.000 | $1.03 \times 10^{-4}$ | 0.359 | $3.50 \times 10^{3}$ |
| 5 | rs344157 | | 5:35034700 | 0.630 | 1.000 | $1.03 \times 10^{-4}$ | 0.359 | $3.50 \times 10^{3}$ |
| 6 | rs40200 | | 5:35045745 | 0.219 | 0.805 | $5.80 \times 10^{-7}$ | $2.17 \times 10^{-3}$ | $3.74 \times 10^{3}$ |
| 7 | rs2291702 | rs11744796 | 5:34993215 | 0.178 | 0.796 | $7.21 \times 10^{-9}$ | $6.68 \times 10^{-6}$ | $9.26 \times 10^{2}$ |
| 8 | | rs16899972 | 5:34998778 | 0.211 | 0.561 | | $3.96 \times 10^{-5}$ | $5.50 \times 10^{3}$ |
| 9 | | rs344512 | 5:35027754 | 0.605 | 1.000 | | $1.60 \times 10^{-5}$ | $2.21 \times 10^{3}$ |
| 10 | | rs465974 | 5:35031048 | 0.615 | 1.000 | | $1.31 \times 10^{-5}$ | $1.82 \times 10^{3}$ |
| 11 | | rs344157 | 5:35034700 | 0.630 | 1.000 | | $1.31 \times 10^{-5}$ | $1.82 \times 10^{3}$ |
| 12 | | rs40200 | 5:35045745 | 0.219 | 0.805 | | $2.47 \times 10^{-5}$ | $3.42 \times 10^{3}$ |

**Supplementary Fig. 15. Functional implication of rs2291702 in AGXT2 expression regulation.** (A) Regional plots of the AGXT2 locus displaying association from no-NAFLD group (upper panel) and NAFLD (lower panel). Color represents $R^2$ between rs2291702 and nearby SNPs. Blue lines denote recombination rate collected from 1000 Genomes Project (Phase 3). (B) ChIP-seq (CTCF and RAD21) tracks within the AGXT2 locus, acquired from ENCODE. The arrow shows direction of AGXT2 transcription. (C,D) Putative enhancer activity of AGXT2 eSNPs assayed by luciferase assay. Luciferase activity between reference and alternative alleles in normal status (C) and in oleic acid/palmitate-treated steatosis model (D). The alternative allele showed reduced transcriptional activity relative to the reference allele, consistently recapitulating the reduced gene expression observed with the alternative allele in human livers. (E) Conditional analysis of rs2291702 and neighboring SNPs (1 to 6, indicated as triangles in (A)). rs2291702 is correlated not only with nearby SNPs but also with SNPs in different

LD blocks near the 3’-end of the main isoform, suggesting that rs2291702 comprises a part of an active enhancer and may interact with distant factors. *. $P < 0.05$, **. $P < 0.01$; independent t-test.

### Supplementary Tables

**Supplementary Table 1. Summary statistics of the participants in the discovery cohort (*n* = 125).**

| | No-NAFLD (*n* = 42) | NAFLD (*n* = 83) | | *P*-value |
|---|---|---|---|---|
| | | NAFL (*n* = 18) | NASH (*n* = 65) | |
| Age, years | 56.7 (12.3) | 50.0 (16.1) | 54.2 (13.5) | 0.29 |
| Male, N (%) | 25 (59.5) | 12 (66.7) | 26 (40.0) | $4.70 \times 10^{-2}$ |
| BMI, kg/m$^2$ | 24.0 (3.4) | 26.7 (3.6) | 28.4 (4.0) | $5.71 \times 10^{-8}$ |
| Fibrosis stage (0-4) | 0.26 (0.5) | 0.78 (0.4) | 1.91 (0.9) | $3.16 \times 10^{-17}$ |
| Significant fibrosis (≥2), N (%) | 1 (2.4) | 0 (0) | 40 (61.5) | $9.44 \times 10^{-12}$ |
| NAFLD activity score | 0.43 (0.59) | 3.17 (1.04) | 5.06 (1.07) | 0.62 |
| SBP, mmHg | 131.0 (16.5) | 126.1 (18.1) | 132.4 (18.5) | 0.56 |
| DBP, mmHg | 79.4 (11.2) | 80.0 (14.1) | 79.8 (12.4) | 1.00 |
| HDL-cholesterol | 46.4 (10.4) | 45.2 (12.8) | 44.6 (13.2) | 0.45 |
| Triglycerides, mg/dL | 123.7 (54.8) | 169 (103.9) | 171.6 (83.2) | $3.46 \times 10^{-3}$ |
| AST, IU/L | 29.9 (18.7) | 36.1 (21.8) | 77.4 (79.3) | $1.52 \times 10^{-11}$ |
| ALT, IU/L | 30.3 (24.8) | 42.5 (22.8) | 92.4 (82.7) | $5.69 \times 10\text{-}^{11}$ |
| GGT, IU/L | 49.4 (53.8) | 48.9 (44.1) | 72.4 (49.5) | $1.12 \times 10^{-4}$ |
| Albumin, g/dL | 4.1 (0.3) | 4.2 (0.2) | 4.1 (0.3) | 0.11 |
| Platelet, ×10$^9$/L | 228.7 (50.7) | 234.7 (63.5) | 221.2 (64.4) | 0.62 |
| HOMA-IR | 2.7 (1.1) | 4.5 (2.6) | 6.7 (5.2) | $1.95 \times 10^{-6}$ |
| Adipo-IR | 5.8 (4.5) | 9.8 (7.2) | 15 (12.3) | $1.10 \times 10^{-6}$ |
| Diabetes, N (%) | 9 (21.4) | 5 (27.8) | 36 (55.4) | $1.13 \times 10^{-3}$ |
| Hypertension, N (%) | 16 (38.1) | 7 (38.9) | 24 (36.9) | 0.99 |

Values are given as mean (SD). *P*-values are from the Kruskal-Wallis test or χ2 test comparing no-NAFLD, NAFL, and NASH groups.

**Supplementary Table 2. Summary statistics of the participants in the replication cohort (*n* = 162).**

| | NAFLD (*n* = 162) | | *P*-value |
|---|---|---|---|
| | NAFL (*n* = 114) | NASH (*n* = 48) | |
| Age, years | 47.4 (16.0) | 50.8 (19.4) | 0.29 |
| Male, N (%) | 71 (62.3) | 19 (39.6) | $7.94 \times 10^{-3}$ |
| BMI, kg/m$^2$ | 29.8 (18.3) | 29.1 (3.7) | 0.67 |
| Fibrosis stage (0-4) | 0.84 (0.78) | 1.79 (0.90) | $1.15 \times 10^{-8}$ |
| Significant fibrosis (≥2), N (%) | 13 (11.4) | 27 (56.3) | $4.11 \times 10^{-9}$ |
| NAFLD activity score | 2.57 (1.15) | 4.88 (1.06) | $2.50 \times 10^{-21}$ |
| SBP, mmHg | 131.0 (15) | 133.4 (14.6) | 0.35 |
| DBP, mmHg | 81.5 (13) | 80.3 (11.7) | 0.56 |
| HDL-cholesterol | 44.3 (10.3) | 44.8 (12) | 0.78 |
| Triglycerides, mg/dL | 149.4 (70.6) | 156.6 (73.2) | 0.56 |
| AST, IU/L | 40.6 (40.8) | 63.3 (29.6) | $1.36 \times 10^{-4}$ |
| ALT, IU/L | 57.8 (59) | 89.1 (66.5) | $5.95 \times 10^{-3}$ |
| GGT, IU/L | 50.0 (46.7) | 75.8 (75.2) | $3.11 \times 10^{-2}$ |
| Albumin, g/dL | 4.2 (0.3) | 4.2 (0.3) | 0.77 |
| Platelet, ×10$^9$/L | 250.8 (62.9) | 238.2 (72.2) | 0.30 |
| HOMA-IR | 5.3 (4.8) | 6.6 (4.9) | 0.15 |
| Adipo-IR | 10.9 (9.0) | 12.9 (7.5) | 0.15 |
| Diabetes, N (%) | 32 (28.1) | 16 (33.3) | 0.50 |
| Hypertension, N (%) | 35 (30.7) | 18 (37.5) | 0.40 |

Values are given as mean (SD). *P*-values are from the independent *t*-test or $\chi 2$ test comparing no-NAFLD, NAFL and NASH groups.

**Supplementary Table 5. Summary statistics of NAFLD participants in the discovery cohort (*n* = 83) by rs2291702 genotype.**

| | CC (*n* = 49) | CT (*n* = 27) | TT (*n* = 7) | *P*-value |
|---|---|---|---|---|
| Age, years | 54.5 (14.1) | 51.6 (14.4) | 51.9 (14.6) | 0.69 |
| Male, N (%) | 22 (44.9) | 14 (51.9) | 2 (28.6) | 0.53 |
| BMI, kg/m$^2$ | 28.4 (4.2) | 28.3 (3.3) | 24.4 (2.8) | $4.31 \times 10^{-2}$ |
| Fibrosis stage (0-4) | 1.69 (0.94) | 1.63 (1.01) | 1.57 (1.27) | $7.76 \times 10^{-3}$ |
| Significant fibrosis (≥ 2), N (%) | 24 (49.0) | 13 (48.1) | 3 (42.9) | 0.96 |
| NAFLD activity score | 4.82 (1.3) | 4.48 (1.34) | 4.14 (1.35) | $8.53 \times 10^{-3}$ |
| SBP, mmHg | 130.6 (20.6) | 133.3 (13.9) | 124.0 (18.9) | 0.26 |
| DBP, mmHg | 79.5 (13.5) | 79.5 (12.3) | 83.7 (9.4) | $9.30 \times 10^{-3}$ |
| HDL-cholesterol | 44.8 (14) | 44.9 (11.3) | 44.0 (14.1) | 0.06 |
| Triglycerides, mg/dL | 168.9 (84.5) | 176.4 (104.4) | 164.6 (35.8) | 0.30 |
| AST, IU/L | 78.6 (87.9) | 54.4 (35.2) | 39.4 (25.3) | $1.57 \times 10^{-2}$ |
| ALT, IU/L | 85.3 (84.9) | 80.6 (64.8) | 46.4 (35.1) | $1.82 \times 10^{-4}$ |
| GGT, IU/L | 66.0 (43.4) | 69.3 (58.8) | 61.3 (51.6) | 0.88 |
| Albumin, g/dL | 4.1 (0.3) | 4.2 (0.3) | 4.1 (0.3) | $6.96 \times 10^{-4}$ |
| Platelet, ×10$^9$/L | 218.0 (60.7) | 237.3 (69.9) | 220.4 (66.1) | $1.88 \times 10^{-4}$ |
| HOMA-IR | 6.5 (5) | 5.7 (3.8) | 5.5 (7.3) | 0.14 |
| Adipo-IR | 12.6 (10.8) | 16.7 (13.2) | 11.4 (7) | 0.74 |
| Diabetes, N (%) | 23 (46.9) | 15 (55.6) | 3 (42.9) | 0.72 |
| Hypertension, N (%) | 18 (36.7) | 11 (40.7) | 2 (28.6) | 0.83 |

Values are given as mean (SD). *P*-values are from the Kruskal-Wallis test or $\chi 2$ test comparing rs2291702 CC, CT, and TT carriers after adjusting for age, sex and BMI.

**Supplementary Table 6. Summary statistics of NAFLD participants in the replication cohort (*n* = 162) by rs2291702 genotype.**

| | CC (*n* = 72) | CT (*n* = 74) | TT (*n* = 16) | *P*-value |
|---|---|---|---|---|
| Age, years | 47.4 (16.7) | 49.4 (17.5) | 48.1 (17.3) | 0.79 |
| Male, N (%) | 42 (58.3) | 38 (51.4) | 10 (62.5) | 0.59 |
| BMI, kg/m$^2$ | 28.7 (3.9) | 28.0 (3.6) | 29.3 (3.0) | 0.22 |
| Fibrosis stage (0-4) | 1.03 (0.84) | 1.16 (0.99) | 1.19 (1.05) | $7.63 \times 10^{-6}$ |
| Significant fibrosis (≥2), N (%) | 14 (19.4) | 19 (25.7) | 6 (37.5) | 0.28 |
| NAFLD activity score | 3.32 (1.40) | 3.18 (1.58) | 3.31 (1.99) | $1.97 \times 10^{-8}$ |
| SBP, mmHg | 133.9 (16.8) | 129.2 (12.3) | 133.5 (15.8) | $3.83 \times 10^{-2}$ |
| DBP, mmHg | 83.4 (12.2) | 78.4 (12.3) | 83.5 (14.2) | $2.70 \times 10^{-3}$ |
| HDL-cholesterol | 42.4 (8.5) | 45.6 (10.9) | 48.4 (17.2) | $2.81 \times 10^{-3}$ |
| Triglycerides, mg/dL | 159.6 (72) | 145.8 (69.2) | 142.0 (77.9) | 0.05 |
| AST, IU/L | 44.8 (26.9) | 50.3 (50) | 45.3 (28) | $3.26 \times 10^{-3}$ |
| ALT, IU/L | 70.5 (63.4) | 61.6 (58) | 76.5 (81.2) | $7.51 \times 10^{-15}$ |
| GGT, IU/L | 61.9 (62.5) | 50.2 (42.8) | 72.4 (87.5) | 0.41 |
| Albumin, g/dL | 4.2 (0.3) | 4.2 (0.3) | 4.2 (0.3) | $2.96 \times 10^{-12}$ |
| Platelet, ×10$^9$/L | 254.8 (70.3) | 238.5 (63.7) | 251.9 (51.6) | $2.20 \times 10^{-8}$ |
| HOMA-IR | 6.3 (6) | 5.5 (3.9) | 4.8 (3.1) | 0.18 |
| Adipo-IR | 12.1 (9.9) | 10.9 (7.2) | 11.5 (8.3) | $3.56 \times 10^{-3}$ |
| Diabetes, N (%) | 19 (26.4) | 27 (36.5) | 2 (12.5) | 0.12 |
| Hypertension, N (%) | 30 (41.7) | 17 (23.0) | 6 (37.5) | 0.05 |

Values are given as mean (SD). *P*-values are from the Kruskal-Wallis test or χ2 test comparing rs2291702 CC, CT, and TT carriers after adjusting for age, sex and BMI.

**Supplementary Table 7.** List of NAFLD-eQTL genes that are also previously associated with NAFLD.

| Gene | β (NAFLD) | β (No-NAFLD) | Previous NAFLD association | References |
|---|---|---|---|---|
| *GLI2* | -0.73 | 0.08 | Heterozygous knockout mice develop more fatty liver when exposed to fatty liver-inducing diets. | Guillen-Sacoto *et al. J Hepatol*. 2017 |
| *SAV1* | -0.70 | -0.05 | Deletion in the liver accelerates the development of NAFLD and liver cancer in mice. | Jeong *et al. J Clin Invest*. 2018 |
| *CD5L* | 0.67 | 0.10 | Serum levels of CD5L is significantly correlated with the stage of liver fibrosis. | Bárcena *et al. EBioMedicine*. 2019 |
| *NR5A2* | -0.60 | -0.05 | Deficient mice fed with high-fat diet displayed macrovesicular steatosis, liver injury, and glucose intolerance, | Miranda *et al. JCI Insight.* 2018 |
| *SLC25A1* | -0.70 | 0.05 | The gene inhibition prevents steatohepatitis, reduces inflammatory macrophage infiltration in the liver and adipose tissue, and mitigates obesity induced by a high-fat diet in mice. | Tan *et al. Cell Death Differ.* 2020 |
| *MLKL* | -0.75 | 0.00 | Inhibition showed protective effects of NASH by decreasing hepatic fat synthesis and chemokine ligand expressions. | Saeed *et al. J Gastroenterol Hepatol.* 2019 |
| *CROT* | 0.68 | -0.08 | Palmitate treatment significantly increases the gene expression via hepatocellular fatty acid oxidation in mouse primary hepatocytes. | Zhao *et al. Sci Rep.* 2015 |
| *VWF* | -0.61 | -0.05 | The gene deficiency reduces the progression of liver fibrosis in mice. | Joshi *et al. Toxicol Appl Pharmacol.* 2017 |
| *HSPA1A* | 0.63 | 0.05 | The gene knockdown decreases fat accumulation in NAFLD, | Zhang *et al. Lipids Health Dis*. 2018 |
| *GPD1* | -0.73 | -0.02 | Mutation causes hepatomegaly, steatohepatitis, and hypertriglyceridemia in early infancy. | Joshi *et al. Eur J Hum Genet.* 2014 |
| *PDGFC* | -0.69 | -0.06 | Both transient and stable expression resulted in the development of liver fibrosis | Campbell *et al. Proc Natl Acad Sci U S* |

| | | | | |
|---|---|---|---|---|
| | | | consisting of the deposition of collagen in a pericellular and perivenular pattern that resembles human alcoholic and nonalcoholic fatty liver disease. | *A.* 2005 |
| *SERPINA6* | 0.55 | 0.01 | Involved in corticosteroid-binding globulin deficiency, which includes fatty liver phenotype. | Moisan and Castanon *Front Endocrinol*. 2016 |
| *LILRB4* | 0.62 | 0.03 | The hepatocyte-specific knockout exacerbated high-fat diet-induced insulin resistance, glucose metabolic imbalance, hepatic lipid accumulation, and systematic inflammation in mice. | Lu *et al.* *Hepatology*. 2018 |

**Supplementary Table 8. Primer sequences used in plasmid construction**

| Probe ID | Orientation | Sequence |
|---|---|---|
| Region 1 | F | TGTGGTAAAATCGATAAGGATCTGCCACTAGAGAGATGACCA |
| | R | CAAGGGCATCGGTCGAAACAGACCATAGCAATGCCT |
| Region 2 | F | TGTGGTAAAATCGATAAGGATCCAGTTTTGCTTCCTCTGGCT |
| | R | CAAGGGCATCGGTCGAGAGGAAGCCACTGTCGAGAT |
| Region 3 | F | TGTGGTAAAATCGATAAGGATCCAGTGGGTAAGGGCATCTGA |
| | R | CAAGGGCATCGGTCGAGAGACCACTTGAGGACACCA |
| Region 4 | F | TGTGGTAAAATCGATAAGGATCGTCCCTACGTTTGTCAAGCC |
| | R | CAAGGGCATCGGTCGACAGAGGTGAATGCGAATGATGT |
| Region 5 | F | TGTGGTAAAATCGATAAGGATCAGAACTCCAATCTTAGGACCTTC |
| | R | CAAGGGCATCGGTCGAGCTTACACTGATGATTAGGAGCA |
| Region 6 | F | TGTGGTAAAATCGATAAGGATCTGAGCCACTGTTCACCAAGA |
| | R | CAAGGGCATCGGTCGACTGGAGGAAGGGCTATGGAA |
